# Transcriptomic dynamics of petal development in the one-day flower species, Japanese morning glory (*Ipomoea nil*)

**DOI:** 10.1101/2024.08.29.610218

**Authors:** Soya Nakagawa, Atsushi Hoshino, Kazuyo Ito, Hiroyo Nishide, Katsuhiro Shiratake, Atsushi J. Nagano, Yasubumi Sakakibara

## Abstract

Various aspects of Japanese morning glory *(Ipomoea nil*) petals, such as color, pattern, shape, flower opening time, and senescence, have been extensively studied. To facilitate such studies, transcriptome data were collected from flower petals at 3-h intervals over 3.5 days; the data was collected from 72 h before and 12 h after flower opening, accounting for 29 timepoints. Data analysis revealed substantial transcriptomic changes before and after flower opening. The expression patterns of cell division marker genes indicated that cell division practically stops at approximately 48 h before flower opening. Furthermore, the increased expression of genes encoding transporters for sugars, amino acids, nucleic acids, and autophagy-related genes was observed after flower opening, indicating the translocation of nutrients from senescing petal cells to other developing tissues. Correlations were found between the temporal expression patterns of the three transcriptional regulators and expression patterns of different sets of structural genes within the anthocyanin biosynthesis pathway, indicating differential reliance on each regulator for the activation of specific structural genes. Furthermore, clock genes were identified. Three copies of the clock gene *ELF3* did not exhibit circadian rhythms, potentially allowing *I. nil* to adapt to high-latitude regions. The temporal transcriptome data and interactive database (https://ipomoeanil.nibb.ac.jp/fpkm/) offer valuable insights into gene expression dynamics, periodicity, and correlations and provide a crucial resource for further research on *I. nil* and other plant species.

## INTRODUCTION

Petals primarily attract pollinators and thus facilitate the pollination success (Nicolson and Wright, 2017). Many plant species have evolved diverse petal colors, patterns, shapes, and scents to achieve this goal (Schiestl, 2015; Moyroud and Glover, 2017; Wozniak and Sicard, 2018; Trunschke *et al*., 2021; Fairnie *et al*., 2022). Some species have developed mechanisms that synchronize blooming with the active periods of their pollinators (van Doorn and van Meeteren, 2003; van Doorn and Kamdee, 2014). Furthermore, flowers have captivated humans since ancient times for their beauty. Breeding efforts have enhanced petal traits in many species, with molecular breeding even producing blue roses (Katsumoto *et al*., 2007; Chandler and Sanchez, 2012). Studies on petals have advanced through primary research focused on understanding the biosynthesis of specialized metabolites responsible for color and scent, as well as the development of cells, tissues, and organs responsible for diverse shapes. Applied research aims at improving these traits for new cultivars, for which whole genome sequences and transcriptomes have become indispensable.

The molecular biology of petals has been studied for over 30 years in model plants such as snapdragon (*Antirrhinum majus*), petunia *(Petunia hybrida)*, and Japanese morning glory (*Ipomoea nil*), which have rich collections of transposon-induced mutants (Schwarz-Sommer *et al*., 2003; Gerats and Vandenbussche, 2005; Morita and Hoshino, 2018; Strazzer *et al*., 2023). Among these, *I. nil*, an annual vine native to tropical America, has been extensively cultivated in Japan as a summer ornamental plant. With a research history spanning over a century, the whole genome sequence for *I*. *nil* was published in 2016 (Hoshino *et al*., 2016a). Furthermore, efficient genome-editing methods have also been established (Watanabe *et al*., 2017). Numerous mutations affecting petal color and shape have been selected from a horticultural perspective, with over 3,000 lines maintained by the National BioResource Project.

*I. nil* petals have been extensively used as a model for studying anthocyanin pigmentation (Morita and Hoshino, 2018). Wild-type *I. nil* produces blue flowers through the accumulation of an anthocyanin known as Heavenly Blue Anthocyanin in the vacuoles (Lu *et al*., 1992). Researchers have isolated the structural genes encoding enzymes involved in anthocyanin biosynthesis and regulatory genes that activate their transcription from flower color mutants (Inagaki *et al*., 1994; Hoshino *et al*., 2001; Hoshino *et al*., 2003; Morita *et al*., 2005; Hoshino *et al*., 2009; Morita *et al*., 2015). Regulatory genes involved in anthocyanin biosynthesis are conserved across species. They encode R2R3-MYB, bHLH, and WDR repeat proteins that form the MBW complex (Quattrocchio *et al*., 2006; Xu *et al*., 2015). Studies on pale-colored mutants led to the discovery of the *EFP* gene, which encodes a chalcone isomerase-like protein that enhances the efficiency of flavonoid synthesis (Morita *et al*., 2014; Waki *et al*., 2020). Furthermore, the cation–proton exchanger PURPLE/InNHX is transiently expressed just before flower opening, increasing vacuolar pH and contributing to the blue coloration of petals (Fukada-Tanaka *et al*., 2000; Yamaguchi *et al*., 2001).

*I. nil* is a 1-day flowering plant, with petals opening synchronously in the morning and wilting by the same evening. The day of flowering can be easily predicted based on the petal size approximately 4 days before flower opening. This characteristic is essential in studying petal growth and senescence. Physiological studies using the Violet line showed that petals fully open 10 h after sunset at 25°C. However, nocturnal light inhibits petal opening, and this inhibition is suppressed by low temperatures (Kaihara and Takimoto, 1979, 1981). The NAC-type transcription factor EPHEMERAL1 (EPH1) controls active petal senescence; thus, suppressing its function delays senescence, allowing flowers to remain open longer (Shibuya *et al*., 2014; Shibuya *et al*., 2018). Studies on the effects of petal morphology using mutants have also explored petal dorsoventrality and glandular trichomes (Iwasaki and Nitasaka, 2006; Shimoki *et al*., 2021). Furthermore, *I. nil* has been studied as a model for photoperiodic flowering (Imamura, 1967). Research on clock genes involved in photoperiodic flowering has confirmed that orthologs of *Arabidopsis* (*A. thaliana*) *GI*, *LHY*, *FKF1*, and *TOC1* exhibit circadian rhythms in leaves (Higuchi *et al*., 2011, Hayama *et al*., 2018).

Transcriptome analyses have been conducted at various developmental stages to understand petal traits. For instance, in *Rosa chinensis*, transcriptomes were obtained at four stages, and the expression of genes related to anthocyanins was compared across these stages (Han *et al*., 2017). In petunia, transcriptomes of petals were sampled at two stages around flower opening, both in the morning and evening, to identify candidate transcription factors that repress scent (Shor *et al*., 2023). These studies typically obtained transcriptomes from two to four stages. However, to analyze genes involved in circadian rhythms or gene co-expression, covering a broader range of developmental stages and conducting large-scale, high-resolution temporal transcriptome analyses at shorter intervals are essential. Such analyses are expected to enhance our understanding of processes, such as the circadian control of floral scent emission timing in petunia (Fenske *et al*., 2015; Fenske and Imaizumi, 2016) and flower opening in *I. nil* (Kaihara and Takimoto, 1979).

Large-scale, high-resolution temporal transcriptome analyses have been conducted in tomato (*Solanum lycopersicum*) leaves grown in plant factories. Analysis of transcriptomes sampled every 2 hours over 2 days identified periodically expressed genes involved in the biosynthesis of salicylic acid, abscisic acid, ethylene, and jasmonic acid (Tanigaki *et al*., 2015). Analysis of transcriptomes from field-grown *Arabidopsis halleri* leaves were sampled every 2 hours over 2 days during equinoxes and solstices revealed clock genes with seasonally varying expression amplitudes (Nagano *et al*., 2019). This study also involved weekly sampling over 2 years, to identify genes with seasonal expression variations. Although such large-scale transcriptome analyses are expensive, methods such as low-cost and easy RNA-seq (Lasy-seq; Kamitani *et al*., 2019) can reduce costs by enabling the inexpensive creation of numerous sequencing libraries.

This study conducted a large-scale, high-resolution temporal transcriptome analysis of *I. nil* petals to understand petal traits comprehensively. Transcriptome samples were collected every 3 hours over 3.5 days, from 72 h before to 12 h after flower opening, covering a total of 29 timepoints. This period encompassed the stages from the closed bud to fully open petals and then to fully closed petals, during which the fresh weight of petals increased approximately tenfold. Library preparation was performed using the Lasy-seq method to minimize the cost per sample.

## RESULTS

### Temporal transcriptome analysis of petals

Large-scale temporal analysis transcriptomes from *I. nil* petals was conducted. The onset of the light period was conveniently defined as the time of flower petal opening when nearly all petals were fully open (Figure S1). To objectively evaluate flower opening, the relative corolla area was measured, which yielded a value of 0.9087 at the time of flower opening (Figure S2).

Petal sampling was performed at 3-hour intervals from 72 h before to 12 h after flower opening, including 29 timepoints. Each sample had three replicates. Petal fresh weight and length were measured (Figure S3a), and total RNA was extracted. The epidermal cell areas on limb, ray, and tube surfaces were measured at −48, −24, and 0 h from another set of plants grown under the same conditions as those from which total RNA was extracted (Figure S3b).

A total of 87 RNA samples were used to prepare sequencing libraries using the Lasy-seq method (Kamitani *et al*., 2019), yielding an average of 8.3 million reads per sample (Table S1). The reads were mapped to two reference transcriptomes (mRNA) and two datasets of predicted long non-coding RNA (lncRNA) sequences (Table 1). One of the predicted lncRNA datasets was generated using a total RNA pool obtained from the 87 samples, resulting in the prediction of 5,364 new lncRNAs. The reads mapped to 73% of the mRNA sequences predicted by Hoshino *et al*. (2016a), 70% and 4% of the mRNA and lncRNA sequences predicted by GenBank (Asagao_1.1, NCBI RefSeq assembly: GCF_001879475.1), respectively, and 7% of the newly predicted lncRNA sequences. The reads mapped to 19,622 mRNA sequences predicted by Hoshino *et al*. (2016a), 17,864 mRNA and 3,045 lncRNA sequences predicted by GenBank, respectively, and 4,623 newly identified lncRNA sequences, capturing their temporal expression changes in petals.

**Table 1.**
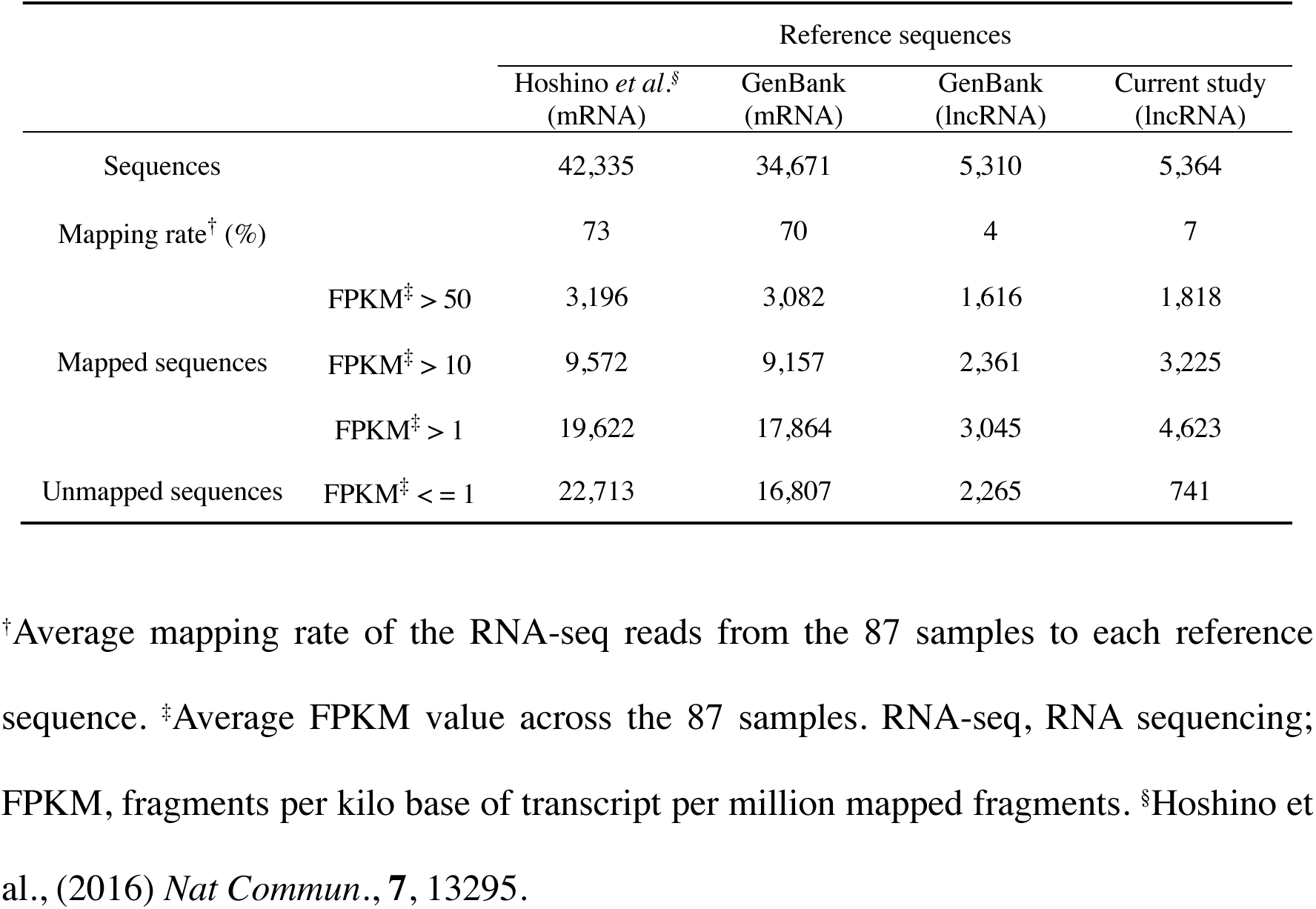
RNA-seq mapping statistics.

An interactive web-based database was constructed to visualize the obtained temporal transcriptomes (Figure 1). This database documented the temporal expression patterns of *PURPLE/InNHX1* and *EPH1* in the petals of *I*. *nil* (Figures 1 and S4). *PURPLE/InNHX1* expression was previously examined by northern hybridization at 12-h intervals from 36 h before to the hour of flower opening (Yamaguchi *et al*., 2001). *EPH1* expression was previously analyzed by reverse transcription quantitative polymerase chain reaction at 4-h intervals up to 32 h after flower opening (Shibuya *et al*., 2014). The changes in the expression of these two genes in the constructed database were consistent with the previously reported expression patterns.

**Figure 1.**
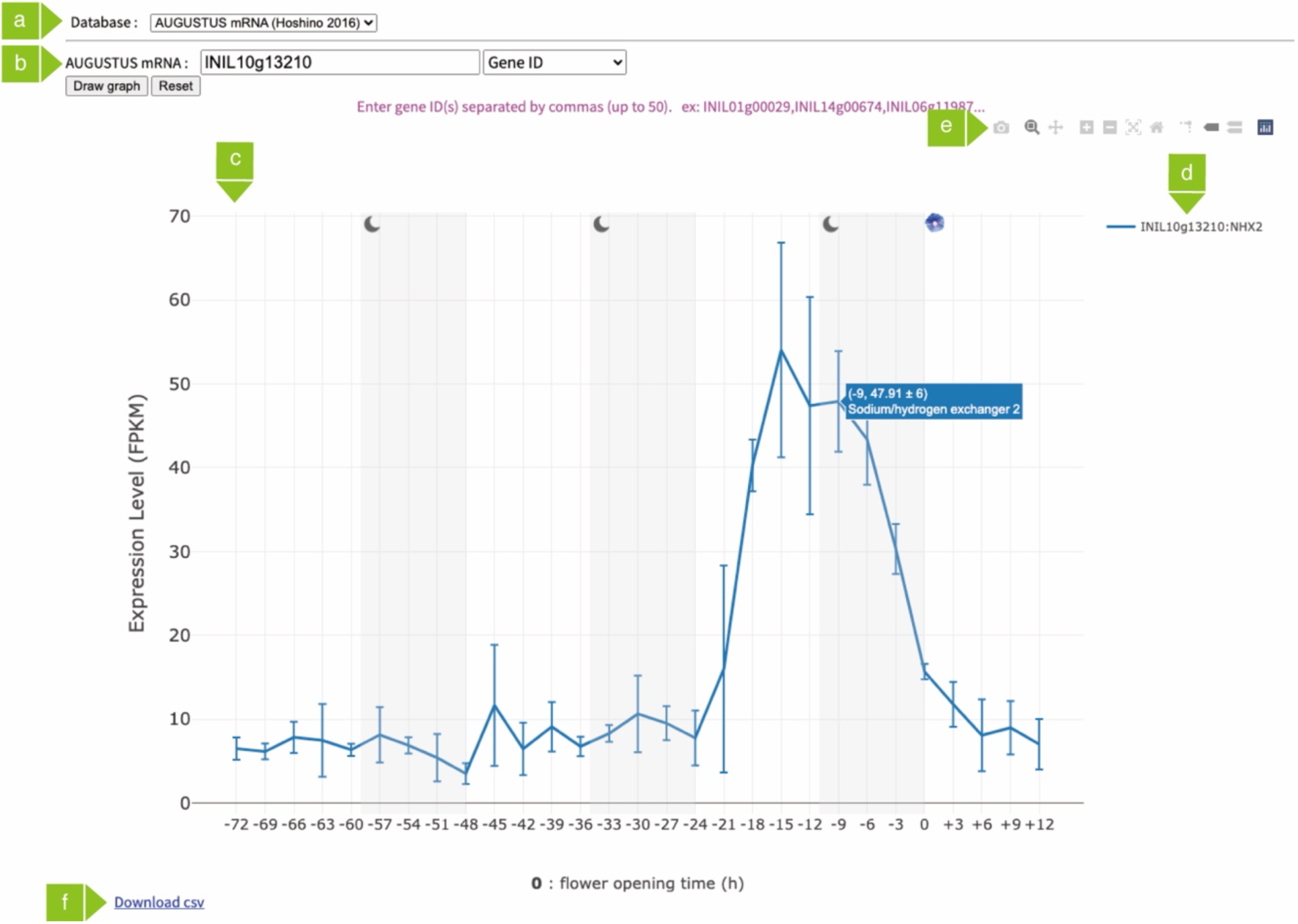
Interface of the temporal transcriptome database for *Ipomoea nil* petals Screen displaying the temporal expression pattern of the *PURPLE/InNHX1* gene (INIL10g13210). (a) The dropdown menu is used to select the database. The expression level databases, which are generated by mapping RNA sequencing reads to four transcript datasets—including mRNA sequences predicted by Hoshino *et al*. (2016a) using AUGUSTUS (Stanke and Waack, 2003); mRNA sequences predicted by GenBank; and lncRNA sequences predicted by GenBank and newly identified in this study—are selectable. (b) A box for entering keywords and a dropdown menu to select the keyword type can be seen. Results are displayed by clicking the “Draw graph” button. (c) Graph area showing the temporal gene expression patterns. The x-axis represents the time from flower opening (0), and the y-axis shows the mean ± standard deviation fragments per kilo base of transcript per million mapped fragments (FPKM) values at each time point (n = 3). The white and gray backgrounds indicate light and dark photoperiods, respectively. Hovering the cursor over the graph reveals the FPKM values and gene descriptions. (d) List of the genes searched. Clicking on a gene hides the graph for that gene, while double-clicking hides the graphs for unselected genes. (e) Buttons are used to pan and zoom the graph and export it as a PNG file. (f) Clicking “Download CSV” allows the graph data to be downloaded as a CSV file.

### Gene ontology analysis

Gene ontology (GO) enrichment analysis was performed to elucidate the biological functions and behaviors of highly expressed genes based on transcriptome data. At each time point, 2,117 genes were selected, representing the top 5% of fragments per kilo base of transcript per million mapped fragments (FPKM) values. Furthermore, the substantially enriched GO terms among these genes were identified. The comprehensive results are presented in Table S2. The eight GO terms with the lowest p-values were highlighted, and the percentage of gene IDs associated with each GO term was calculated (Figure 2).

**Figure 2.**
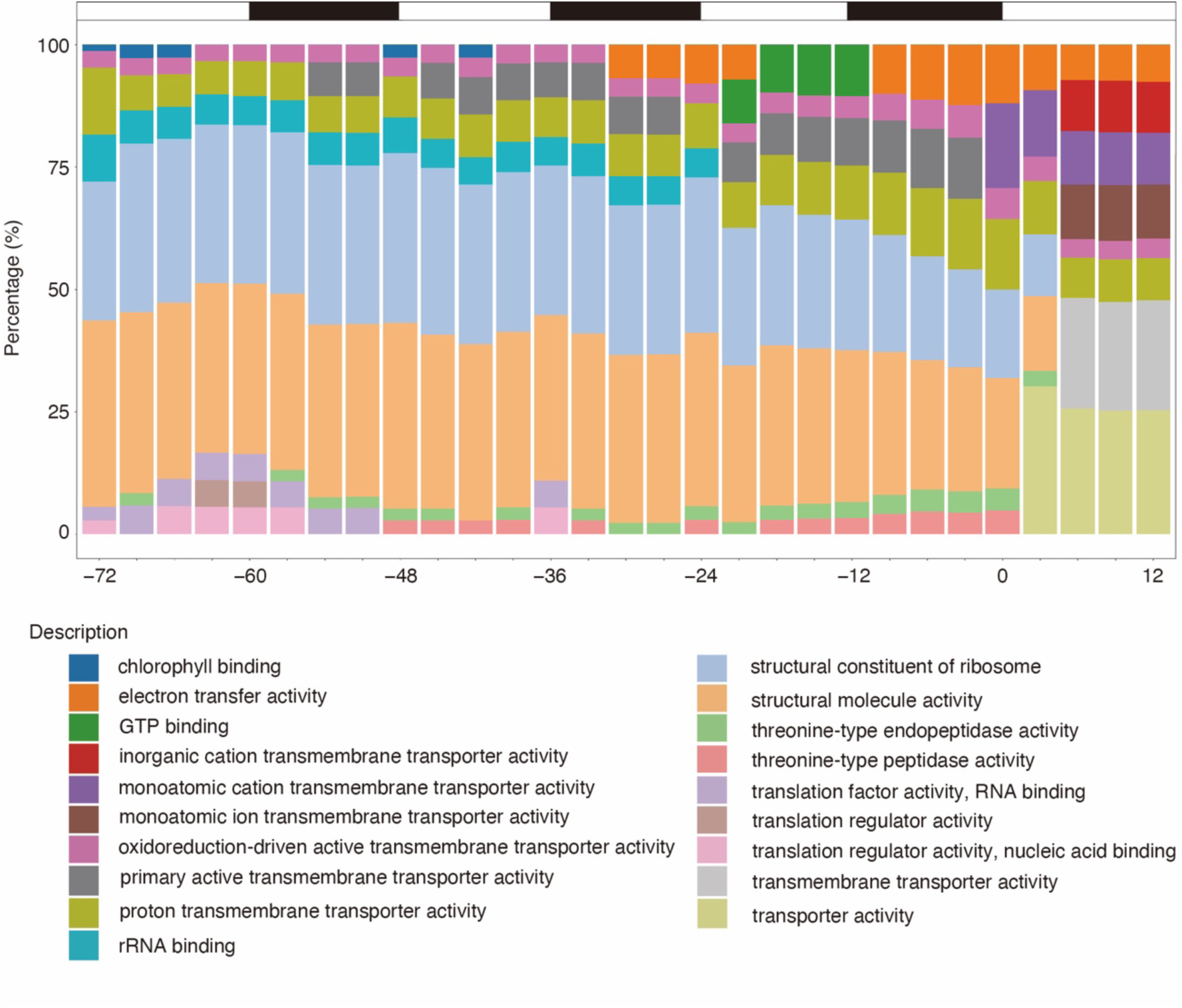
Results of gene ontology (GO) enrichment analysis. The eight GO terms classified under “Molecular Function” with the lowest p-values were selected. The y-axis shows the percentage of gene IDs associated with each term. The x-axis indicates the time from flower opening (0). The bar at the top of the figure indicates the photoperiod. White and black bars indicate light and dark conditions, respectively.

Between −72 and −42 h during the light period, “chlorophyll binding,” a GO term associated with photosynthesis, was particularly prominent. From −72 to 0 h, GO terms related to mRNA translation were prevalent. However, during petal wilting, 6–12 h after flower opening, the prevalence of mRNA translation-related GO terms decreased. In contrast, transporter-related GO terms emerged as seven of the top eight. This shift reflects a substantial change in dominant GO terms before and after flower opening.

The annotations of genes associated with the top seven transporter-related GO terms at 12 h after flower opening were examined to characterize the transporters expressed during petal wilting. The analysis revealed that 13, 19, 9, and 14 genes were involved in the transport of sugars, nitrogen (including amino acids), phosphates (including nucleotides), and inorganic ions, respectively (Table S3). Visualization of the gene expression levels using a heatmap indicated that, with one or two exceptions in each of the four substrate-classified groups, transporter gene expression increased following flower opening (Figure S5). Notably, INIL03g42103, a homolog of *SWEET12*, which is a gene implicated in sucrose transport from mesophyll cells to phloem in *Arabidopsis* (Chen *et al*., 2012), obtained an FPKM value of approximately 11,400 at 12 h after flower opening, representing 1.14% of total transcripts.

### Evaluation and identification of internal control genes

In comparative gene expression analysis, internal control genes with relatively high and stable expression levels are commonly used for normalization. However, the expression stability of internal control genes in *I. nil* petals has not been previously established. Therefore, the expression variations of seven genes previously used as internal controls in petals were visualized using the newly created database (Figure S6a). Furthermore, the coefficient of variation (CV) was calculated as an index of expression stability for these genes (Table S4). The gene with the smallest CV value (0.219) was INIL09g35921 encoding a mitochondrial ATP synthase γ-subunit. To identify more suitable internal control genes beyond the seven originally analyzed, the CV values of all genes were calculated, and the top five genes with average FPKM values >100 and the lowest CV values were visualized in the database (Figure S6b). The gene with the smallest CV value at 0.072 was the *Arabidopsis DEFICIENS* homolog encoding a MADS-box protein (INIL15g27881; Table S4), making it a suitable internal control gene.

### Analysis of periodic genes and circadian clock genes

To identify genes exhibiting circadian rhythms, transcriptome data were analyzed using MetaCycle. At a threshold of BH.Q < 0.05, 805 genes displaying circadian rhythms were identified (Figure 3a and Table S5). *PnC401* (Sage-Ono *et al*., 1998), which exhibits circadian rhythms in cotyledons, also exhibited circadian rhythm in petals (INIL05g23942 in Table S5). GO enrichment analysis of these genes identified many terms related to photosynthesis, light response, and circadian rhythms (Figure 3b and Table S6). No genes were identified with ultradian rhythms of 12-h periodicity at the BH.Q < 0.05 threshold (Table S7). Furthermore, lncRNAs exhibiting circadian rhythms were identified using MetaCycle (Tables S8–S9). Of the lncRNAs exhibiting circadian rhythms identified, 32 exhibiting circadian rhythms at the BH.Q < 0.05 threshold were registered in GenBank, whereas 58 were newfound.

**Figure 3.**
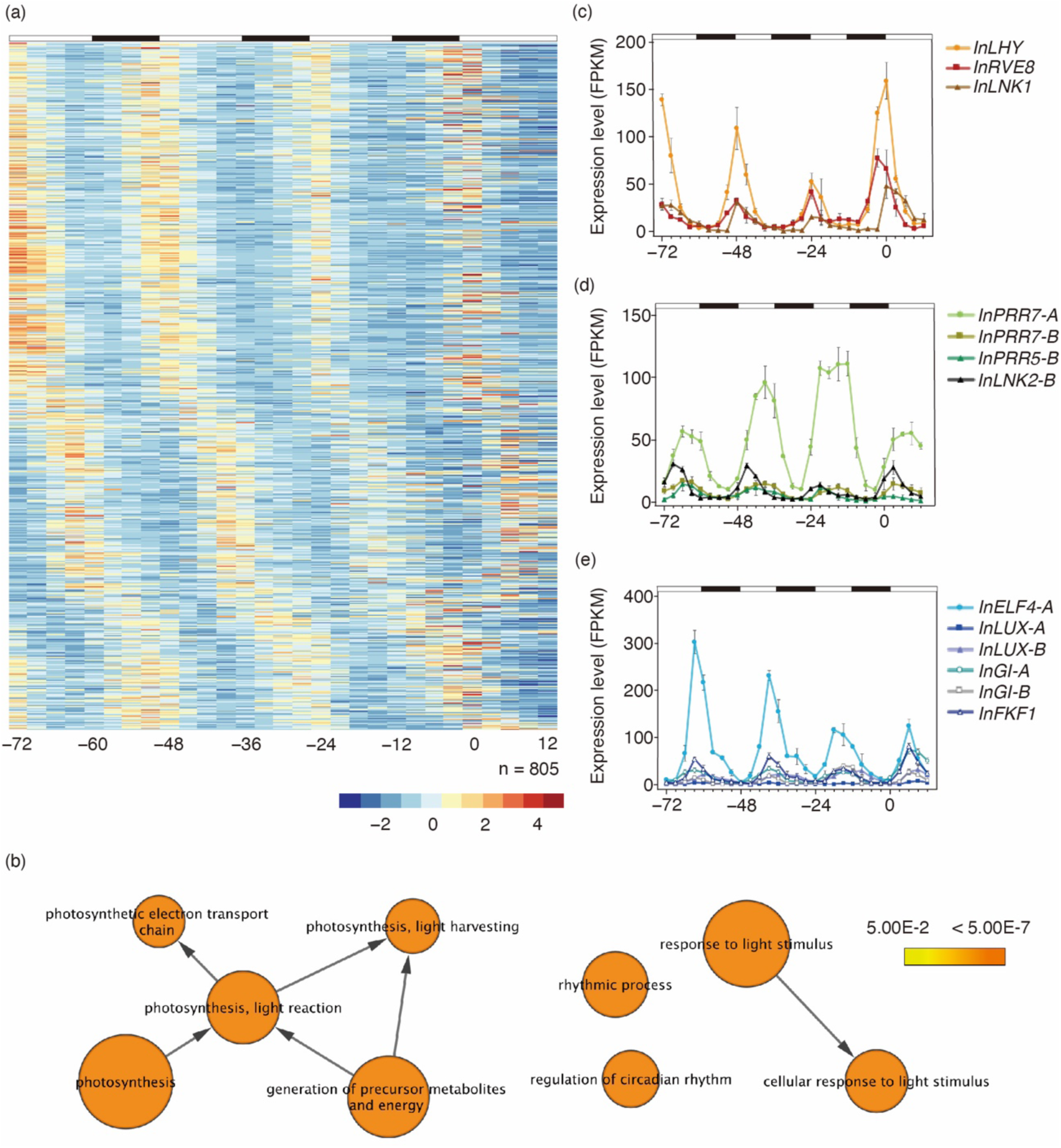
Expression analysis of genes with circadian rhythms, including clock genes. (a) Heatmap of 805 genes exhibiting circadian rhythms. The fragments per kilo base of transcript per million mapped fragments (FPKM) values for each gene were normalized using Z-scores to express relative expression levels. The color scale represents Z-scores, with a gradient from red to blue indicating high to low expression. The bar at the top denotes the photoperiod, with white and black bars representing light and dark conditions, respectively. (b) GO enrichment analysis. Nine enriched GO terms with the lowest p-values were selected and visualized as nodes in a network based on the hierarchical structure of GO terms. The size of each node indicates the number of genes included, and the color represents p-values according to the color scale. (c–e) Expression patterns of clock genes exhibiting a 24-h circadian rhythm with a BH.Q value < 0.0001. The x-axis represents the time from flower opening (0), while the y-axis shows the mean ± standard deviation FPKM values at each time point (n = 3). The bar at the top indicates the photoperiod, with white and black bars representing light and dark conditions, respectively. The expression patterns of clock genes are shown with peaks at (c) dawn, (d) during the daytime, (e) and in the evening.

Furthermore, clock genes exhibiting circadian rhythms in petals were identified from the list of 805 genes(Table S5). Clock genes were explored further by analyzing protein sequences from *I. nil* and six other plants (*A. thaliana*, *Oryza sativa*, *Vitis vinifera*, *Fragaria vesca*, *Lotus japonicus*, *S. lycopersicum*) using OrthoFinder to create ortholog groups (Tables S10 and S11). Thirty-two ortholog groups containing well-known *Arabidopsis* clock genes were identified (Table S12). Of these, 7 groups contained no *I. nil* orthologs, while the remaining 25 ortholog groups comprised 49 orthologs in *I. nil*. Notably, although orthologs of *CCA1*, which encodes a core component of the circadian clock, were not found in the six plants other than *Arabidopsis*, orthologs of *LHY*, a clock gene highly homologous to *CCA1*, were identified. *I. nil* clock genes (*PnGI*, *PnLHY*, *RVE1*, *PRR7*, *PnFKF1*, and *PnTOC1*) previously characterized were also identified as orthologs of the corresponding *Arabidopsis* genes using OrthoFinder (Table S13; Higuchi *et al*., 2011; Shinozaki *et al*., 2014; Hayama *et al*., 2018). Of the 49 orthologs, 44 were expressed at average FPKM levels >1 in petals, with 18 exhibiting circadian rhythms at the BH.Q < 0.05 threshold (Figures 3c–3e and Table S12). Among *InGI-A*, *InLHY*, *InFKF1*, and *InTOC1*, which exhibit circadian rhythms in leaves (Higuchi *et al*., 2011, Hayama *et al*., 2018), *InTOC1* did not exhibit circadian rhythms in petals (Table S12). Additionally, although *ELF3*, a core oscillator component in *Arabidopsis*, exhibits circadian rhythm (Hicks *et al*., 2001, Huang and Nusinow, 2016), none of the three orthologs identified in *I. nil* (*InELF3-A*, *InELF3-B*, *InELF3-C*) exhibited circadian rhythms (Table S12).

### Expression analysis of genes related to cell division and the cell cycle

Petal growth is driven by increased cell number and size, with cell number playing a more substantial role during the early stages of growth than cell size (Krizek and Anderson, 2013). Therefore, the expression of genes regulating the cell cycle and cell division, which increase the cell number, was investigated using transcriptome data. The orthologs of 14 *Arabidopsis* gene families involved in the regulation of the cell cycle and cell division in *I. nil* were determined using OrthoFinder (Table S14). Orthologs were identified for all gene families, totaling 47 genes. A heatmap was generated from the transcriptome data to examine the expression changes of these 47 genes. Based on the heatmap, the expression patterns of the genes were classified into five clusters (Clusters A–E; Figure 4a).

**Figure 4.**
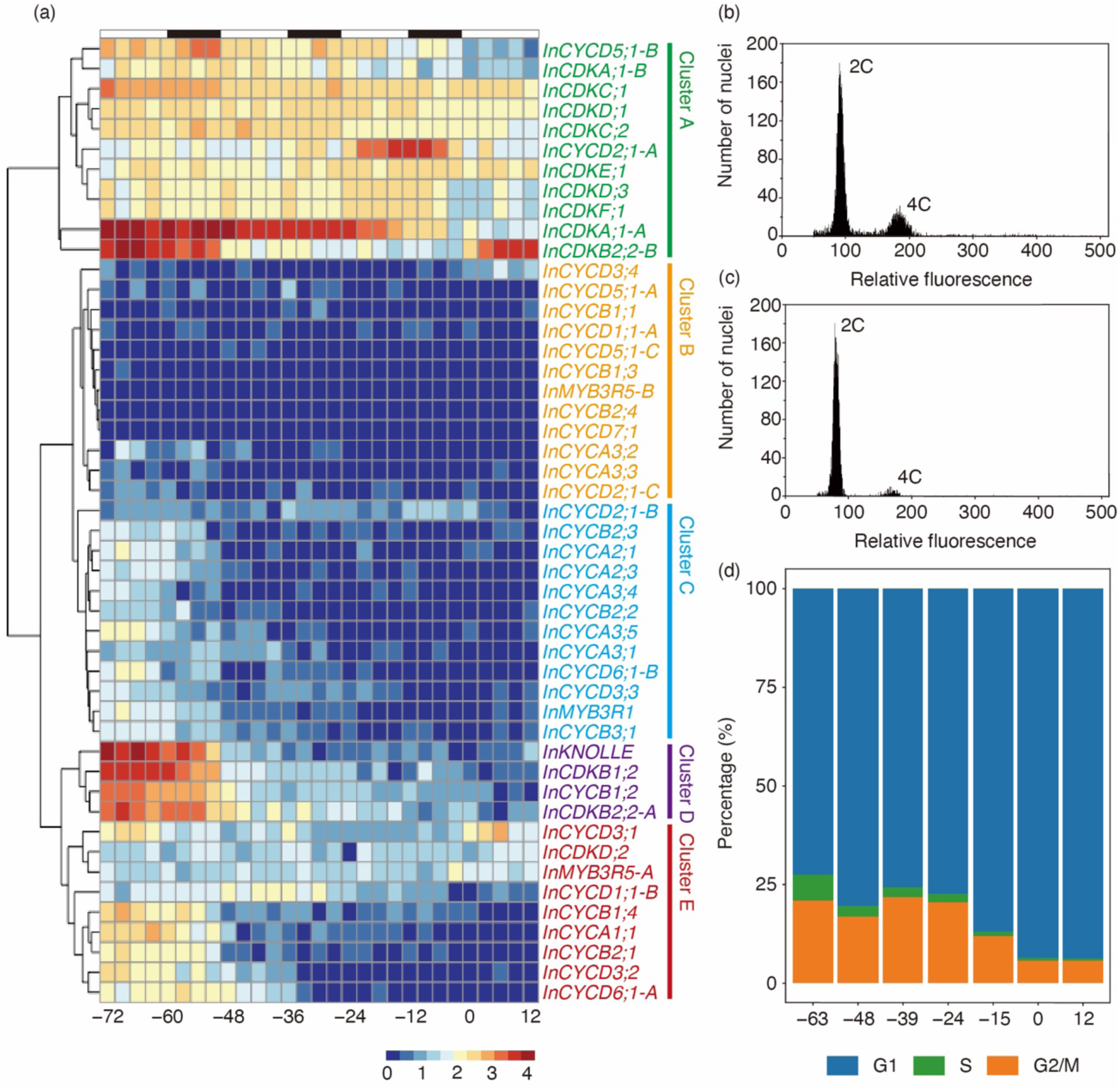
Analysis of cell division and cell cycle of petal cells (a) Heatmap analysis of the temporal expression levels of genes involved in cell division and the cell cycle. Expression values are expressed as log_10_ (FPKM+1). The x-axis represents the time from flower opening (0), with the bar at the top indicating the photoperiod. White and black bars represent light and dark conditions, respectively. (b–d) Cell cycle analysis. The DNA content of the nuclei of petal cells was measured by flow cytometry (b) 63 h before and (c) 12 h after flower opening. (d) Percentages of cells in the G1, S, and G2/M phases. The x-axis indicates the time from flower opening (0), and the y-axis shows the proportion of cells in each cell cycle phase.

Four genes in Cluster D (*InKNOLLE, InCDKB1;2, InCDKB2;2-A,* and *InCYCB1;2*), whose *Arabidopsis* orthologs served as markers for cell division, exhibited decreasing expression levels from −72 h, reaching a basal level approximately −48 h before flower opening (Figure 4a). In *Arabidopsis*, *KNOLLE* encodes a cytokinesis-specific syntaxin, which accumulates exclusively during the M phase (Lukowitz *et al*., 1996; Lauber *et al*., 1997). *CDKB* and *CYCB* are crucial for G2/M phase progression (Stals and Inzé, 2001; Gutierrez, 2009; Qi and Zhang, 2020). These observations indicate that cell division in *I. nil* petals remains active until approximately −48 h before flower opening.

The cell cycle of petal cells was analyzed through flow cytometry at seven different time points, from 63 h before to 12 h after flower opening (Figures 4b–4d). The proportion of cells in the S phase decreased to 6.54% at −63 h before flower opening, with the most substantial reduction occurring between −63 and −48 h before opening (down to 2.8%). Conversely, the proportion of cells in the G2/M phase decreased markedly from 24 h before flower opening until the time of opening. After flower opening, no substantial changes were observed in the proportions of cells at each cell cycle stage. Further, no distinct peaks beyond 8C were detected, aligning with previous findings on the cell cycle in leaves (Hoshino *et al*., 2016a).

### Analysis of the expression of anthocyanin biosynthesis genes

Using transcriptome data, we visualized the expression levels (Figure S7) and changes in expression (Figure 5a) of genes involved in anthocyanin biosynthesis (Table S15) through heatmaps. These expression changes were grouped into four distinct clusters (Clusters I–IV), with three regulatory genes (*InMYB1*, *InbHLH2*, and *InWDR1*) distributed among the different clusters (Figure 5a). *InMYB1* demonstrated a peak in expression approximately −15 h before flower opening, forming a cluster (Cluster IV) with five structural genes that exhibited relatively low expression levels until −27 h before flower opening. Cluster IV included four genes (*InCHS-D*, *InEFP*, *InF3H-C*, and *InDFR-B*) involved in the upstream steps of the anthocyanin biosynthesis pathway before DFR and one gene (*InGST1*) coding for an enzyme that acts downstream. *InbHLH2* also peaked in expression around −15 h before flower opening and retained more than half of its peak expression level at −27 h before flower opening. This gene formed Cluster III with five structural genes, which encode biosynthetic enzymes that function downstream of CHI in the anthocyanin biosynthesis pathway (*InCHI*, *InF3H-A*, *InANS*, *In3GT*, and *In3GGT*). *InWDR1* exhibited relatively low expression levels around −15 h before flower opening and formed a cluster (Cluster II) with *InCHS-E* and *InF3′H*, which peaked at −48 and −45 h before flower opening, respectively. *InCHS-E* exhibited 24-h periodic expression (INIL14g35461 in Table S5), whereas *InWDR1* did not. Conversely, *InF3′H* displayed periodic increases in expression every 24 h at −72, −48, and −24 h before flower opening, similar to *InCHS-E*, but did not increase at the time of flower opening.

**Figure 5.**
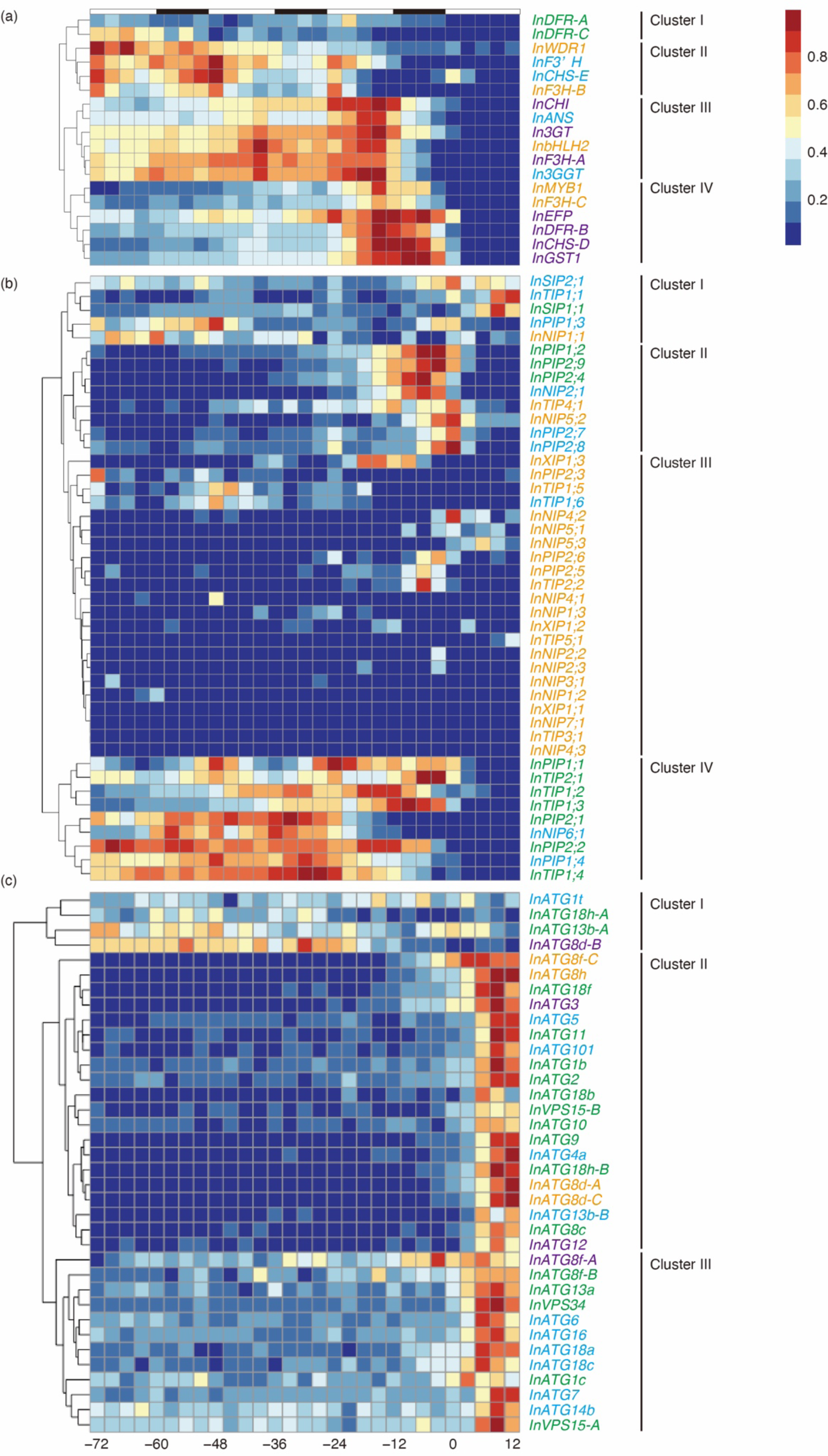
Expression patterns of genes related to anthocyanin biosynthesis, aquaporins, and autophagy Fragments per kilo base of transcript per million mapped fragments (FPKM) values for each gene were min–max normalized to a range of 0–1, with colors on the scale representing values from blue to red. The x-axis represents the time from flower opening (0), and the bar at the top denotes the photoperiod, with white and black bars indicating light and dark conditions, respectively. The colors of the gene names correspond to clusters obtained from heatmaps created using non-normalized FPKM values (Figures S7–S9). The lists of anthocyanin biosynthesis and autophagy-related genes are provided in Tables S15 and S18, respectively. Aquaporin genes were referenced from Inden *et al*. (2023) and are listed in Table S17. Expression patterns of (a) anthocyanin biosynthesis genes, (b) aquaporin genes, and (c) autophagy-related genes.

The correlation between the expression changes of the three regulatory genes and the structural genes was further confirmed by Pearson correlation coefficient analysis (Table S16). However, the correlation coefficients of *InF3′H* with *InbHLH2* and *InWDR1* were equivalent. In *Arabidopsis*, *RVE8*/*LCL5*, which encodes a circadian clock component, is involved in the 24-h periodic expression of structural genes (Perez-Garcia *et al*., 2015). Among the *I. nil RVE* genes, two exhibited circadian rhythms in the petals, *InRVE8*, the ortholog of *Arabidopsis RVE8*, and *InRVE1-A* (Table S12). However, the expression of *InRVE8* and *InRVE1-A* were not correlated with the expression of any structural genes, including *InCHS-E* and *InF3′H* (Table S16). These results indicate that different mechanisms in *Arabidopsis* and *I*. *nil* drive the 24-h periodic expression of structural genes.

### Expression analysis of aquaporin genes

A genome-wide analysis of *I. nil* identified 44 aquaporin genes whose functions were predicted (Inden *et al*., 2023). Heatmaps were used to visualize both expression levels (Figure S8) and expression changes (Figure 5b) to explore the expression patterns of these aquaporin genes. The expression changes were grouped into four clusters (Clusters I–IV) (Figure 5b). Cluster I comprised genes that were expressed during the early stages of petal development or at and after flower opening. Cluster II included genes that showed high expression starting from 12 h before flower opening. Cluster III contained genes with consistently low expression levels, while Cluster IV included genes with elevated expression from 72 h before to the time of flower opening.

The fresh weight and cell volume of petals increase rapidly, starting at 24 h before flower opening, with a significant acceleration increase within the final 12 h (Figure S3). Aquaporin genes in Cluster II, whose expression increased in tandem with this increase in weight and volume—particularly those encoding aquaporins that function in the cell membrane (*InPIP1;2*, *InPIP2;4*, and *InPIP2;9*)—may facilitate the rapid water influx into petal cells, a process essential for flower opening. The peak FPKM value of *InPIP1;2* reached approximately 15,500, accounting for 1.55% of the total transcript abundance. These findings underscore the crucial role of *InPIP1;2* in flower opening.

Aquaporin genes in Cluster IV may play a critical role in water and small molecule transport, thereby substantially contributing to petal development given their high gene expression beginning at −72 h before flower opening. Five of the 44 aquaporin genes exhibited circadian rhythms at a BH.Q value < 0.05 (Table S17). Of these five genes, *InPIP1;1* and *InTIP2;1*, which are classified in Cluster IV, showed higher expression levels than the other three genes (Figure S8). The physiological functions of these aquaporins and their relationship with circadian rhythms warrant further investigation.

### Expression analysis of autophagy-related genes

In *I. nil* petals, autophagy is triggered by the progression of senescence, resulting in elevated expression levels of two autophagy-related genes, *InATG8* and *InATG4* (Shibuya *et al*., 2009a;Shibuya *et al*., 2009b; Shibuya *et al*., 2011). In plants, the core autophagy mechanism involves 19 proteins, including ATG1–ATG14, ATG16, ATG18, ATG101, VPS15, and VPS34 (Avila-Ospina *et al*., 2014; Liu *et al*., 2018; Norizuki *et al*., 2019). Therefore, the transcriptome was analyzed to determine whether autophagy-related genes and *InATG8* and *InATG4* exhibit a coordinated increase in expression during petal senescence.

Orthologs of the 19 autophagy-related gene families in *Arabidopsis* were identified in *I. nil* using OrthoFinder, which identified orthologs for all 19 families (Tables S13 and S18). The *ATG1, ATG8, ATG13, ATG18*, and *VPS15* families had multiple copies (3, 8, 3, 6, and 2, respectively). In contrast, the remaining gene families had a single copy, resulting in 36 genes encoding proteins involved in the core autophagy mechanism in the *I. nil* genome. Heatmaps were generated to examine the expression patterns of the 45 autophagy-related genes identified (Figures 5c and S9). Notably, 41 genes exhibited peak expression levels 6 h after flower opening (Figure 5c). The four genes that peaked earlier belonged to the multi-copy *ATG1, ATG8, ATG13*, and *ATG18* families, indicating that the expression levels of all 19 gene families increased in a coordinated manner following flower opening. Furthermore, four genes coding for ATG8, which plays a central role in autophagosome formation (*InATG8d-A, InATG8d-C, InATG8f-C,* and *InATG8h*), formed a cluster with genes that were particularly highly expressed (Cluster D in Figure S9).

## DISCUSSION

A large-scale, high-resolution temporal transcriptome of *I. nil* petals was collected and analyzed across 3.5 days. The analysis was performed from 72 h before to 12 h after flower opening at 3-h intervals, totaling 29 timepoints. Transcriptome collection from petals at various developmental stages has also been conducted in other plants such as rose, pineapple, lily, daffodil, and petunia (Han *et al*., 2017; He *et al*., 2020; Wang *et al*., 2020; Yin *et al*., 2020; Shor *et al*., 2023). Based on morphological characteristics, these studies commonly classified petals into two to four stages. In contrast, the present study classified petals into 29 timepoint stages based on the time relative to flower opening. This classification was made possible by the characteristic simultaneous flowering of *I. nil* at a specific time (Figures S1–S3). This approach allows for a more detailed examination of petal development through temporal transcriptome changes. Moreover, collecting transcriptomes every 3 h facilitated the comprehensive identification of gene groups with approximately 24-h periodicity in petals (Figure 3a and Table S5). Furthermore, genes with minimal expression variation over the 3.5 days that could serve as internal controls were identified (Figure S6 and Table S4).

In this study, 87 transcriptomes were obtained by collecting three samples at each of the 29 timepoints. The Lazy-seq method (Kamitani *et al*., 2019) enabled library preparation and sequencing at approximately one-fourth the cost of conventional methods. Specifically, transcriptomes of approximately 20,000 genes were obtained at approximately 20 cents per gene.

### Petal growth by cell division and enlargement

Four key insights were obtained by analyzing the temporal transcriptome dynamics. First, petal growth was linked to the increase in both cell division and cell size. Growth in *I. nil* petals resulted from the increase in cell number through division and cell volume. The length and weight of petals continuously increased from −72 to −3 h before flower opening (Figure S3). The synchronized decrease in the expression levels of four cell division marker genes (*InKNOLLE*, *InCDKB1;2*, *InCDK2;2*, and *InCYCB1;2*) and the significant reduction in the proportion of cells in the S phase approximately −48 h before flower opening indicate that cell division nearly ceased at this time (Figures 4a–4d). Therefore, petal growth up to −48 h before flower opening is likely driven by both cell division and cell enlargement, while growth from −48 to −3 h before flower opening is primarily due to cell enlargement. These findings are consistent with the fact that early-stage plant tissue growth is more significantly influenced by an increase in cell number than by cell volume expansion (Gonzalez *et al*., 2012; Krizek and Anderson, 2013; Hepworth and Lenhard, 2014).

Furthermore, the cell volume increase rapidly, starting at −24 h before flower opening, as indicated by changes in petal length, weight, and cell surface area (Figure S3). Aquaporin genes classified in Cluster II—which showed increased expression during this period (especially *InPIP1;2*, accounting for 1.55% of the total transcript abundance from −6 to −3 h before flower opening)—are likely responsible for the water influx required for cell volume expansion (Figure 5b). Indeed, aquaporin inhibitors suppress flower opening in *I. nil* (Katsuhara *et al*., 2008). Co-expression analysis of petal length, weight, and transcriptome data obtained in this study will provide insights into the gene networks underlying cell enlargement and volume expansion.

### Senescence and nutrient translocation

The second key finding involves petal senescence and nutrient translocation. The transcriptome undergoes substantial changes before and after flower opening, with GO terms related to transporters being enriched after flower opening (Figure 2). The expression levels of transporter genes for sugars, nitrogen (including amino acids), phosphates (including nucleic acids), and inorganic ions (Figure S5), as well as those of autophagy-related genes (Figures 5c and S9), increased during this period. These results indicate increased degradation and transport activities within senescing petal cells. Petal senescence is a type of active programmed cell death (van Doorn and Woltering, 2008). In *I. nil*, the NAC-type transcription factor EPH1 activates this process (Shibuya *et al*., 2014, Shibuya *et al*., 2018). During senescence, the expression of autophagy-related genes, such as *ATG4* and *ATG8*, increases, and autophagosome-like structures are observed (Shibuya *et al*., 2009a; Yamada *et al*., 2009; Shibuya *et al*., 2011). This study found that at least one gene from each of the 19 gene families involved in forming the core autophagy mechanism peaked in expression 6 h after flower opening (Figure 5c). This finding reinforces the conventional understanding that autophagy is induced during petal senescence. Furthermore, these observations indicate a mechanism for transferring and recycling nutrients from dying petal cells to other developing tissues, including the recycling of autophagy-derived degradation products. In senescing petunia petals, phosphorus, nitrogen, and potassium contents decrease, indicating the possible recycling of nutrients from senescing petals (Verlinden, 2003). In senescing leaves, the sugar transporter SWEET is implicated in sucrose translocation (Eom *et al*., 2015). The highly expressed SWEET gene, INIL03g42103, which accounts for 1.14% of the total transcripts at 12 h after flower opening, may play a major role in such translocation processes (Figure S5). Creating and analyzing *I. nil* mutants of transporters and autophagy-related genes that increase in expression after flower opening through genome editing would be a practical approach to investigating this recycling mechanism.

### Anthocyanin biosynthesis

The third key finding pertains to the temporal regulation of anthocyanin biosynthesis genes. The transcriptional activation of structural genes for anthocyanin biosynthesis is controlled by the conserved MBW complex, composed of three transcriptional regulators across species (Quattrocchio *et al*., 2006, Xu *et al*., 2015). In *I. nil*, *InMYB1*, *InbHLH2*, and *InWDR1* are responsible for anthocyanin biosynthesis regulation (Morita *et al*., 2006; Hoshino *et al*., 2016b). *InMYB1* and *InWDR1* mutants produce white flowers, with anthocyanin pigmentation lost either specifically in the petals or throughout the entire plant (Morita *et al*., 2006). Furthermore, *InbHLH2* mutants in close relatives of *I. nil*, *I. purpurea*, and *I. tricolor* produce flowers with paler pigmentation compared with wild-type plants (Park *et al*., 2004; Park *et al*., 2007). In the petals of these *Ipomoea* mutants, the accumulation of the mRNA of these structural genes generally decreases, with few exceptions (Park *et al*., 2004; Morita *et al*., 2006; Park *et al*., 2007). This study demonstrated through heatmap and correlation analyses that the expression changes of each regulatory gene correlated with those of different groups of structural genes (Figure 5a and Table S16). Specifically, the regulatory gene *InMYB1* formed a cluster with the structural genes *InCHS-D*, *InEFP*, *InF3H-C*, *InDFR-B*, and *InGST1*; *InbHLH2* clustered with *InCHI*, *InF3H-A*, *InANS*, *In3GT*, and *In3GGT*; and *InWDR1* formed a cluster with *InCHS-E* and *InF3’H*. Therefore, the transcriptional activation of individual structural genes depends differently on each of the three transcriptional regulators within the same complex. The molecular mechanisms behind this differential transcriptional regulation remain unclear. The ratio of transcriptional regulators binding to the promoters of each structural gene may change over time, and protein-level analyses will be required to elucidate this process.

### Adaptation of *I. nil* to high-latitude regions

The fourth key finding pertains to the identification of genes associated with circadian rhythms, revealing an unexpected outcome: none of the three copies of the clock gene *InELF3* exhibited circadian rhythmicity (Table S12). The Tokyo Kokei Standard line, which was utilized in this study, was initially collected in Japan and is presumably well-adapted to the local environmental conditions, particularly the photoperiod. *I. nil*, which originates from tropical America (Imamura *et al*., 1966; Austin *et al*., 2001; Muñoz-Rodríguez *et al*., 2019), may have adapted to the photoperiod of Japan, a region situated at a higher latitude, by losing the circadian rhythm in *InELF3* expression.

In *Arabidopsis*, *ELF3* is a core component of the circadian clock, with both ELF3 mRNA and protein levels exhibiting circadian rhythms (Hicks *et al*., 2001; Huang and Nusinow, 2016). *ELF3* regulates flowering in response to light and temperature stimuli. In contrast, *elf3* mutants exhibit early flowering phenotypes regardless of photoperiod conditions (Zagotta *et al*., 1996; Huang and Nusinow, 2016). In barley (*Hordeum vulgare*) and legumes (*Pisum sativum* and *Lens culinaris*), loss-of-function mutations in *ELF3* orthologs have been selected to confer early flowering and enable cultivation at higher latitudes (Faure *et al*., 2012; Weller *et al*., 2012). In barley *elf3* mutants, the amplitude of expression of clock genes encoding core oscillators declines while the expression of the florigen gene *HvFT1* increases, thereby leading to early flowering (Faure *et al*., 2012). Similarly, *I. nil* may have been selected for *ELF3* mutations that do not exhibit circadian rhythms during its northward expansion. A comparative analysis of the *ELF3* genes between lines from tropical American strains and lines from Japan could verify this hypothesis.

The large-scale, high-resolution temporal transcriptome analysis performed in this study has unveiled the dynamic nature, periodicity, and interrelationships of gene expression within the petals of *I. nil*. This research offers insights that extend beyond petal development, encompassing a wide array of physiological processes. The resulting database will substantially advance studies on pigmentation, pattern formation, petal morphology, flower opening times, and senescence in these short-lived flowers. Furthermore, the database provides a valuable resource for investigating flower petals in other plant species.

## EXPERIMENTAL PROCEDURES

### Plant materials

Seeds were obtained by self-pollinating an individual Tokyo Kokei Standard line plant, which had been used for whole genome sequencing (Hoshino *et al*., 2016a). Four plants were grown from the harvested seeds in commercial potting soil using 15-cm-diameter plastic pots and incubated in an LPH-411SP incubator (Nihon Ika Kikai, Osaka, Japan). The incubator was set to a 13-h light/11-h dark photoperiod at 25°C and 50% humidity. Petal sampling was performed every 3 h, and during the dark period, green light was used to prevent exposure to other wavelengths. The flower opening time was defined as the beginning of the light period (8:00 AM). Sampling was performed from 72 h before flower opening (8:00 AM, 3 days prior) to 12 h after flower opening (8:00 PM on the day of opening) at 29 timepoints. Petals were weighed in the light, photographed with a Nikon Z50 digital camera (Nikon, Tokyo, Japan), frozen in liquid nitrogen, and stored at −85°C. ImageJ Fiji was used to measure petal length on digital camera images (Schindelin *et al*., 2012). Each sample consisted of 3–7 petals.

### Petals and petal cells analysis

The Tokyo Kokei Standard line grown under the conditions described above was used to measure the corolla area. Time-lapse photography was performed every 30 min from 15 h before flower opening to 12 h after flower opening using a Nikon D5100 digital camera. The camera was positioned to face the petals, and images were captured at a resolution of 300 dpi. The camera’s built-in flash was used for photographing in the dark. The corolla area was measured from the images using software that calculates pixel values for the colored regions of the petals (Shinozaki *et al*., 2011) at each time point.

The cell cycle of petal cells was analyzed using a flow cytometer. Petals were cut into 5-mm squares, and nuclei were extracted and stained with 4′,6-diamidino-2-phenylindole using a Quantum Stain NA UV 2 Kit (Partec, Münster, Germany) according to the manufacturer’s instructions. The stained nuclei were analyzed using a CyFlow SL flow cytometer and Partec FloMax software (both from Partec, Münster, Germany), and the proportion of nuclei in each cell cycle phase was calculated.

The area of petal cells was measured from images obtained with a TM4000Plus tabletop scanning electron microscope (Hitachi High-Tech, Tokyo, Japan) using ImageJ Fiji (Schindelin *et al*., 2012). The limb, ray, and tube areas of 100 cells from three flowers collected at each time point were measured.

### RNA extraction and sequencing

Total RNA was extracted from the petals using the Maxwell RSC RNA Plant Kit (Promega, Madison, WI, USA), according to the manufacturer’s protocol with modifications. First, 70–200 mg of frozen pulverized petal cells were placed in 500 µL homogenization buffer with 10 µL thioglycerol and then ground with a mortar and pestle at room temperature. The mixture was centrifuged, and the supernatant was isolated and placed in lysis buffer. RNA was quantified using the Qubit RNA BR Assay Kit and a Qubit fluorometer (both from Thermo Fisher Scientific, Waltham, MA, USA). RNA was used to prepare sequencing libraries following the Lasy-seq protocol (Kamitani *et al*., 2019). Paired-end 150-bp sequencing was performed on a HiSeq X Ten platform (Illumina, San Diego, CA, USA) at Clockmics Inc. (Osaka, Japan). The reads were preprocessed using the Clockmics Inc. pipeline and mapped to four sets of *I. nil* transcripts. Gene expression levels were quantified using FPKM. The mRNA sequences predicted by Hoshino *et al*. (2016a), the mRNA and lncRNA sequences predicted by GenBank (Asagao_1.1, NCBI RefSeq assembly: GCF_001879475.1), and newly obtained lncRNA sequences were used for mapping. The mRNA sequences identified by Hoshino *et al*. (2016a) were mapped, including splice variants, with FPKM values summed for each gene. For the mRNA sequences predicted by GenBank, the longest splice variant for each gene was selected for use.

For lncRNA detection, total RNA obtained from the 29 timepoints and used in Lasy-seq was pooled in equal amounts. rRNA was removed, and cDNA was synthesized using random hexamers to create libraries. Eighty-eight million reads were obtained using the BGISEQ-500 platform (BGI, Shenzhen, China) and mapped to the *I. nil* genome using HISAT 2.0.4 (Kim *et al*., 2019). Transcripts were predicted with StringTie 1.0.4 (Pertea *et al*., 2015), and 26,874 transcripts were identified. Coding potential was determined using Coding Potential Calculator (CPC; Kong *et al*., 2007), txCdsPredict (Kent *et al*., 2002), Coding-Non-Coding Index (CNCI; Sun *et al*., 2013), and the Pfam database (Finn *et al*., 2016), and 5,364 novel lncRNAs were predicted. Their expression levels were also quantified as FPKM values.

An interactive online database was developed to determine the FPKM values for each mRNA and lncRNA (available at https://ipomoeanil.nibb.ac.jp/fpkm/). The backend database management was handled using SQLite3 (https://www.sqlite.org/), while the graph rendering was implemented using Plotly.js (https://plotly.com/javascript/).

### Bioinformatics

Gene expression periodicity was analyzed using MetaCycle version 1.2.0 (Wu *et al*., 2016). Genes that exhibited a meta2d BH.Q value < 0.05 were classified as periodic. GO enrichment analysis was performed using BiNGO version 3.0.3 (Maere *et al*., 2005) with the threshold at p-value < 0.05. GO terms were sourced from Hoshino *et al*. (2016a), and the ontology files were downloaded from the GO website (https://geneontology.org/docs/download-ontology/). Orthologs were identified using OrthoFinder version 2.5.5 (Emms and Kelly, 2019) based on protein sequences from seven plant species (Table S10).

Heatmaps of gene expression levels and changes were generated using pheatmap version 1.0.12 (Kolde and Kolde, 2015). Log_10_-transformed FPKM values, with an addition of 1, were used to construct the heatmap of expression levels. For the heatmaps depicting expression changes, genes identified as periodic on MetaCycle were normalized using a Z-score. In contrast, other genes were normalized using min–max normalization across the 87 transcriptomes, with values averaged across the 29 time points. Anthocyanin biosynthesis genes were mapped using NCBI annotations (Asagao_1.1, NCBI RefSeq assembly: GCF_001879475.1), while other genes were mapped using annotations from Hoshino *et al*. (2016a).

### Statistical analysis

Normality was assessed using the Anderson–Darling test, with a significance threshold of p < 0.05. For two-group comparisons where normality was not met, the Brunner–Munzel test was performed using the lawstat package (version 3.6) in R (Wallace *et al*., 2008) at a significance threshold of p < 0.05.

## ACCESSION NUMBERS

The raw sequencing data and newly identified lncRNA sequences have been deposited in the DNA Data Bank of Japan (DDBJ)/BioProject under accession number PRJDB18471. Additionally, the expression data for predicted genes and lncRNAs, including FPKM values, are available in DDBJ/GEA under the accession number E-GEAD-856.

## ACKNOWLEDGMENTS

The authors wish to thank Shizuka Koizumi for her guidance on bioinformatics analysis. Thanks are also extended to Masayoshi Kawaguchi, Mitsuki Yoshimoto, and Masaki Ishikawa for their valuable discussions and to Tomoyo Takeuchi, Saki Kawada, Kiyoko Kuzunishi, and Naoko Koyama for their technical support. This study utilized equipment at the Model Organisms Facility, Data Integration and Analysis Facility, Trans-Omics Facility, and Optics and Imaging Facility, NIBB Trans-Scale Biology Center. JSPS KAKENHI supported part of this study to AH (18H04127, 18K06301, 19H00944, and 21K06239) and SN (23KJ1004).

The authors declare no conflicts of interest.

## SUPPORTING INFORMATION

**Figure S1.** Petals used for RNA extraction

**Figure S2.** Measurement of relative corolla area

**Figure S3.** Temporal changes in the fresh weight and length of petals and measurement of the area of their epidermal cells

**Figure S4.** Expression pattern of the *EPH1* gene

**Figure S5.** Heatmap analysis of transporter genes

**Figure S6.** Expression patterns of internal control genes

**Figure S7.** Heatmap analysis of the temporal expression levels of anthocyanin biosynthesis genes

**Figure S8.** Heatmap analysis of the temporal expression levels of aquaporin genes **Figure S9.** Heatmap analysis of the temporal expression levels of autophagy-related genes

**Table S1.** RNA sequencing statistics for the 29 timepoints with biological triplicates **Table S2.** Gene ontology enrichment analysis for the transcriptomes of the 29 timepoints

**Table S3.** Gene list for the top eight enriched gene ontology terms of the transcriptome at 12 h after flower opening

**Table S4.** Coefficients of variation for internal control genes

**Table S5.** List of genes with approximately 24-h rhythmic expression identified by MetaCycle analysis (BH.Q < 0.05)

**Table S6.** Gene ontology enrichment analysis of genes with approximately 24-h rhythmic expression

**Table S7.** MetaCycle analysis results for approximately 12-h rhythmic expression (p-value < 0.05)

**Table S8.** List of lncRNAs registered in GenBank with approximately 24-h rhythmic expression identified by MetaCycle analysis (BH.Q < 0.05)

**Table S9.** List of newly identified lncRNAs with approximately 24-h rhythmic expression identified by MetaCycle analysis (BH.Q < 0.05)

**Table S10.** List of genomes used for Orthofinder analysis

**Table S11.** Orthologous groups among the seven species detected by Orthofinder

**Table S12.** Orthologous groups of circadian clock-related genes in seven species evaluated

**Table S13.** Correspondence of gene names between previous and current studies

**Table S14.** *Ipomoea nil* orthologs of cell division and cell cycle genes in *Arabidopsis thaliana*

**Table S15.** List of anthocyanin biosynthesis genes in *Ipomoea nil*

**Table S16.** Correlation coefficients for expression patterns between transcriptional regulator genes and structural genes involved in anthocyanin biosynthesis

**Table S17.** List of aquaporin genes in *Ipomoea nil*

**Table S18.** *Ipomoea nil* orthologs of *Arabidopsis thaliana* autophagy-related genes

## Notes

### Competing Interest Statement

The authors have declared no competing interest.

## REFERENCES

Austin, D.F., Kitajima, K., Yoneda, Y. and Qian, L.F. (2001) A putative tropical American plant, *Ipomoea nil* (convolvulaceae), in pre-Columbian Japanese art. Econ. Bot., 55, 515–527.

Avila-Ospina, L., Moison, M., Yoshimoto, K. and Masclaux-Daubresse, C. (2014) Autophagy, plant senescence, and nutrient recycling. J. Exp. Bot., 65, 3799–3811.

Chandler, S.F. and Sanchez, C. (2012) Genetic modification; the development of transgenic ornamental plant varieties. Plant Biotechnol. J., 10, 891–903.

Chen, L.-Q., Qu, X.-Q., Hou, B.-H., Sosso, D., Osorio, S., Fernie, A.R. and Frommer, W.B. (2012) Sucrose efflux mediated by SWEET proteins as a key step for phloem transport. Science, 335, 207–211.

Emms, D.M. and Kelly, S. (2019) OrthoFinder: phylogenetic orthology inference for comparative genomics. Genome Biol., 20.

Eom, J.-S., Chen, L.-Q., Sosso, D., Julius, B.T., Lin, I.W., Qu, X.-Q., Braun, D.M. and Frommer, W.B. (2015) SWEETs, transporters for intracellular and intercellular sugar translocation. Curr. Opin. Plant Biol., 25, 53–62.

Fairnie, A.L.M., Yeo, M.T.S., Gatti, S., Chan, E., Travaglia, V., Walker, J.F. and Moyroud, E. (2022) Eco-Evo-Devo of petal pigmentation patterning. Essays Biochem., 66, 753–768.

Faure, S., Turner, A.S., Gruszka, D., Christodoulou, V., Davis, S.J., von Korff, M. and Laurie, D.A. (2012) Mutation at the circadian clock gene *EARLY MATURITY 8* adapts domesticated barley (*Hordeum vulgare*) to short growing seasons. Proc. Natl. Acad. Sci. USA, 109, 8328–8333.

Fenske, M.P., Hazelton, K.D.H., Hempton, A.K., Shim, J.S., Yamamoto, B.M., Riffell, J.A. and Imaizumi, T. (2015) Circadian clock gene directly regulates the timing of floral scent emission in. Proc. Natl. Acad. Sci. USA, 112, 9775–9780.

Fenske, M.P. and Imaizumi, T. (2016) Circadian rhythms in floral scent emission. Front. Plant Sci., 7, 462.

Finn, R.D., Coggill, P., Eberhardt, R.Y., Eddy, S.R., Mistry, J., Mitchell, A.L., Potter, S.C., Punta, M., Qureshi, M., Sangrador-Vegas, A., Salazar, G.A., Tate, J. and Bateman, A. (2016) The Pfam protein families database: towards a more sustainable future. Nucleic Acids Res., 44, D279–285.

Fukada-Tanaka, S., Inagaki, Y., Yamaguchi, T., Saito, N. and Iida, S. (2000) Colour-enhancing protein in blue petals. Nature, 407, 581.

Gerats, T. and Vandenbussche, M. (2005) A model system comparative for research: *Petunia*. Trends Plant Sci., 10, 251–256.

Gonzalez, N., Vanhaeren, H. and Inzé, D. (2012) Leaf size control: complex coordination of cell division and expansion. Trends Plant Sci., 17, 332–340.

Gutierrez, C. (2009) The Arabidopsis cell division cycle. Arabidopsis Book, 7, e0120.

Han, Y., Wan, H., Cheng, T., Wang, J., Yang, W., Pan, H. and Zhang, Q. (2017) Comparative RNA-seq analysis of transcriptome dynamics during petal development in *Rosa chinensis*. Sci. Rep., 7, 43382.

Hayama, R., Mizoguchi, T. and Coupland, G. (2018) Differential effects of light-to-dark transitions on phase setting in circadian expression among clock-controlled genes in *Pharbitis nil*. Plant Signal Behav., 13, e1473686.

He, Y.S., Xu, M. and Chen, X.J. (2020) De novo transcriptomics analysis of the floral scent of Chinese narcissus. Trop. Plant Biol., 13, 172–188.

Hepworth, J. and Lenhard, M. (2014) Regulation of plant lateral-organ growth by modulating cell number and size. Curr. Opin. Plant Biol., 17, 36–42.

Hicks, K.A., Albertson, T.M. and Wagner, D.R. (2001) *EARLY FLOWERING3* encodes a novel protein that regulates circadian clock function and flowering in Arabidopsis. Plant Cell, 13, 1281–1292.

Higuchi, Y., Sage-Ono, K., Sasaki, R., Ohtsuki, N., Hoshino, A., Iida, S., Kamada, H. and Ono, M. (2011) Constitutive expression of the *GIGANTEA* ortholog affects circadian rhythms and suppresses one-shot induction of flowering in *Pharbitis nil*, a typical short-day plant. Plant Cell Physiol., 52, 638–650.

Hoshino, A., Jayakumar, V., Nitasaka, E., Toyoda, A., Noguchi, H., Itoh, T., Shin-I, T., Minakuchi, Y., Koda, Y., Nagano, A.J., Yasugi, M., Honjo, M.N., Kudoh, H., Seki, M., Kamiya, A., Shiraki, T., Carninci, P., Asamizu, E., Nishide, H., Tanaka, S., Park, K.-I., Morita, Y., Yokoyama, K., Uchiyama, I., Tanaka, Y., Tabata, S., Shinozaki, K., Hayashizaki, Y., Kohara, Y., Suzuki, Y., Sugano, S., Fujiyama, A., Iida, S. and Sakakibara, Y. (2016a) Genome sequence and analysis of the Japanese morning glory *Ipomoea nil*. Nat. Commun., 7, 13295.

Hoshino, A., Johzuka-Hisatomi, Y. and Iida, S. (2001) Gene duplication and mobile genetic elements in the morning glories. Gene, 265, 1–10.

Hoshino, A., Morita, Y., Choi, J.D., Saito, N., Toki, K., Tanaka, Y. and Iida, S. (2003) Spontaneous mutations of the flavonoid 3’-hydroxylase gene conferring reddish flowers in the three morning glory species. Plant Cell Physiol., 44, 990–1001.

Hoshino, A., Park, K.I. and Iida, S. (2009) Identification of *r* mutations conferring white flowers in the Japanese morning glory (*Ipomoea nil*). J. Plant Res., 122, 215–222.

Hoshino, A., Yoneda, Y. and Kuboyama, T. (2016b) A *Stowaway* transposon disrupts the *InWDR1* gene controlling flower and seed coloration in a medicinal cultivar of the Japanese morning glory. Genes Genet. Syst., 91, 37–40.

Huang, H. and Nusinow, D.A. (2016) Into the evening: Complex interactions in the *Arabidopsis* circadian clock. Trends Genet., 32, 674–686.

Imamura, S. ed (1967) Physiology of Flowering in *Pharbitis nil* Tokyo: Japanese Society of Plant Physiologists.

Imamura, S., Muramatsu, M., Kitajo, S. and Takimoto, A. (1966) Varietal difference in photoperiodic behavior of *Pharbitis nil*. Bot. Mag. Tokyo, 79, 714–721.

Inagaki, Y., Hisatomi, Y., Suzuki, T., Kasahara, K. and Iida, S. (1994) Isolation of a *Suppressor-mutator/Enhancer*-like transposable element, *Tpn1*, from Japanese morning glory bearing variegated flowers. Plant Cell, 6, 375–383.

Inden, T., Hoshino, A., Otagaki, S., Matsumoto, S. and Shiratake, K. (2023) Genome-wide analysis of aquaporins in Japanese morning glory (*Ipomoea nil*). Plants, 12, 1511.

Iwasaki, M. and Nitasaka, E. (2006) The *FEATHERED* gene is required for polarity establishment in lateral organs especially flowers of the Japanese morning glory (*Ipomoea nil*). Plant Mol. Biol., 62, 913–925.

Kaihara, S. and Takimoto, A. (1979) Environmental factors controlling the time of flower-opening in *Pharbitis nil*. Plant Cell Physiol., 20, 1659–1666.

Kaihara, S. and Takimoto, A. (1981) Effects of light and temperature on flower-opening of *Pharbitis nil*. Plant Cell Physiol., 22, 215–221.

Kamitani, M., Kashima, M., Tezuka, A. and Nagano, A.J. (2019) Lasy-Seq: a high-throughput library preparation method for RNA-Seq and its application in the analysis of plant responses to fluctuating temperatures. Sci. Rep., 9, 7091.

Katsuhara, M., Hanba, Y.T., Shiratake, K. and Maeshima, M. (2008) Expanding roles of plant aquaporins in plasma membranes and cell organelles. Funct. Plant Biol., 35, 1–14.

Katsumoto, Y., Fukuchi-Mizutani, M., Fukui, Y., Brugliera, F., Holton, T.A., Karan, M., Nakamura, N., Yonekura-Sakakibara, K., Togami, J., Pigeaire, A., Tao, G.Q., Nehra, N.S., Lu, C.Y., Dyson, B.K., Tsuda, S., Ashikari, T., Kusumi, T., Mason, J.G. and Tanaka, Y. (2007) Engineering of the rose flavonoid biosynthetic pathway successfully generated blue-hued flowers accumulating delphinidin. Plant Cell Physiol., 48, 1589–1600.

Kent, W.J., Sugnet, C.W., Furey, T.S., Roskin, K.M., Pringle, T.H., Zahler, A.M. and Haussler, D. (2002) The human genome browser at UCSC. Genome Research, 12, 996–1006.

Kim, D., Paggi, J.M., Park, C., Bennett, C. and Salzberg, S.L. (2019) Graph-based genome alignment and genotyping with HISAT2 and HISAT-genotype. Nat. Biotechnol., 37, 907-+.

Kolde, R. and Kolde, M.R. (2015) Package ‘pheatmap’. R package, 1, 790.

Kong, L., Zhang, Y., Ye, Z.Q., Liu, X.Q., Zhao, S.Q., Wei, L. and Gao, G. (2007) CPC: assess the protein-coding potential of transcripts using sequence features and support vector machine. Nucleic Acids Res., 35, W345–349.

Krizek, B.A. and Anderson, J.T. (2013) Control of flower size. J. Exp. Bot., 64, 1427–1437.

Lauber, M.H., Waizenegger, I., Steinmann, T., Schwarz, H., Mayer, U., Hwang, I., Lukowitz, W. and Jürgens, G. (1997) The *Arabidopsis* KNOLLE protein is a cytokinesis-specific syntaxin. J. Cell Biol., 139, 1485–1493.

Liu, F., Hu, W. and Vierstra, R.D. (2018) The vacuolar protein sorting-38 subunit of the *Arabidopsis* phosphatidylinositol-3-kinase complex plays critical roles in autophagy, endosome sorting, and gravitropism. Front. Plant Sci., 9, 781.

Lu, T.S., Saito, N., Yokoi, M., Shigihara, A. and Honda, T. (1992) Acylated peonidin glycosides in the violet-blue cultivars of *Pharbitis nil*. Phytochemistry, 31, 659–663.

Lukowitz, W., Mayer, U. and Jurgens, G. (1996) Cytokinesis in the Arabidopsis embryo involves the syntaxin-related KNOLLE gene product. Cell, 84, 61–71.

Maere, S., Heymans, K. and Kuiper, M. (2005) BiNGO: a Cytoscape plugin to assess overrepresentation of Gene Ontology categories in Biological Networks. Bioinformatics, 21, 3448–3449.

Morita, Y. and Hoshino, A. (2018) Recent advances in flower color variation and patterning of Japanese morning glory and petunia. Breed Sci, 68, 128–138.

Morita, Y., Hoshino, A., Kikuchi, Y., Okuhara, H., Ono, E., Tanaka, Y., Fukui, Y., Saito, N., Nitasaka, E., Noguchi, H. and Iida, S. (2005) Japanese morning glory *dusky* mutants displaying reddish-brown or purplish-gray flowers are deficient in a novel glycosylation enzyme for anthocyanin biosynthesis, UDP-glucose:anthocyanidin 3-*O*-glucoside-2″-*O*-glucosyltransferase, due to 4-bp insertions in the gene. Plant J., 42, 353–363.

Morita, Y., Ishiguro, K., Tanaka, Y., Iida, S. and Hoshino, A. (2015) Spontaneous mutations of the UDP-glucose:flavonoid 3-*O*-glucosyltransferase gene confers pale- and dull-colored flowers in the Japanese and common morning glories. Planta, 242, 575–587.

Morita, Y., Saitoh, M., Hoshino, A., Nitasaka, E. and Iida, S. (2006) Isolation of cDNAs for R2R3-MYB, bHLH and WDR transcriptional regulators and identification of *c* and *ca* mutations conferring white flowers in the Japanese morning glory. Plant Cell Physiol., 47, 457–470.

Morita, Y., Takagi, K., Fukuchi-Mizutani, M., Ishiguro, K., Tanaka, Y., Nitasaka, E., Nakayama, M., Saito, N., Kagami, T., Hoshino, A. and Iida, S. (2014) A chalcone isomerase-like protein enhances flavonoid production and flower pigmentation. Plant J., 78, 294–304.

Moyroud, E. and Glover, B.J. (2017) The evolution of diverse floral morphologies. Curr. Biol., 27, R941–R951.

Muñoz-Rodríguez, P., Carruthers, T., Wood, J.R.I., Williams, B.R.M., Weitemier, K., Kronmiller, B., Goodwin, Z., Sumadijaya, A., Anglin, N.L., Filer, D., Harris, D., Rausher, M.D., Kelly, S., Liston, A. and Scotland, R.W. (2019) A taxonomic monograph of *Ipomeoa* integrated across phylogenetic scales. Nat. Plants, 5, 1136–1144.

Nagano, A.J., Kawagoe, T., Sugisaka, J., Honjo, M.N., Iwayama, K. and Kudoh, H. (2019) Annual transcriptome dynamics in natural environments reveals plant seasonal adaptation. Nat. Plants, 5, 329–329.

Nicolson, S.W. and Wright, G.A. (2017) Plant-pollinator interactions and threats to pollination: perspectives from the flower to the landscape. Funct. Ecol., 31, 22–25.

Norizuki, T., Kanazawa, T., Minamino, N., Tsukaya, H. and Ueda, T. (2019) *Marchantia polymorpha*, a new model plant for autophagy studies. Front Plant Sci, 10, 935.

Park, K., Choi, J., Hoshino, A., Morita, Y. and Iida, S. (2004) An intragenic tandem duplication in a transcriptional regulatory gene for anthocyanin biosynthesis confers pale-colored flowers and seeds with fine spots in *Ipomoea tricolor*. Plant J., 38, 840–849.

Park, K.I., Ishikawa, N., Morita, Y., Choi, J.D., Hoshino, A. and Iida, S. (2007) A bHLH regulatory gene in the common morning glory, *Ipomoea purpurea*, controls anthocyanin biosynthesis in flowers, proanthocyanidin and phytomelanin pigmentation in seeds, and seed trichome formation. Plant J., 49, 641–654.

Perez-Garcia, P., Ma, Y., Yanovsky, M.J. and Mas, P. (2015) Time-dependent sequestration of RVE8 by LNK proteins shapes the diurnal oscillation of anthocyanin biosynthesis. Proc. Natl. Acad. Sci. USA, 112, 5249–5253.

Pertea, M., Pertea, G.M., Antonescu, C.M., Chang, T.C., Mendell, J.T. and Salzberg, S.L. (2015) StringTie enables improved reconstruction of a transcriptome from RNA-seq reads. Nat. Biotechnol., 33, 290–295.

Qi, F.F. and Zhang, F.X. (2020) Cell cycle regulation in the plant response to stress. Front. Plant Sci., 10, 1765.

Quattrocchio, F., Baudry, A., Lepiniec, L. and Grotewold, E. (2006) The regulation of flavonoid biosynthesis. In The Science of Flavonoids (Grotewold, E. ed. New York: Springer, pp. 97–122.

Sage-Ono, K., Ono, M., Harada, H. and Kamada, H. (1998) Accumulation of a clock-regulated transcript during flower-inductive darkness in *Pharbitis nil*. Plant Physiol, 116, 1479–1485.

Schiestl, F.P. (2015) Ecology and evolution of floral volatile-mediated information transfer in plants. New Phytol., 206, 571–577.

Schindelin, J., Arganda-Carreras, I., Frise, E., Kaynig, V., Longair, M., Pietzsch, T., Preibisch, S., Rueden, C., Saalfeld, S., Schmid, B., Tinevez, J.Y., White, D.J., Hartenstein, V., Eliceiri, K., Tomancak, P. and Cardona, A. (2012) Fiji: an open-source platform for biological-image analysis. Nat. Methods, 9, 676–682.

Schwarz-Sommer, Z., Davies, B. and Hudson, A. (2003) An everlasting pioneer: the story of *Antirrhinum majus* research. Nat. Rev. Genet., 4, 657–666.

Shibuya, K., Shimizu, K., Niki, T. and Ichimura, K. (2014) Identification of a NAC transcription factor, EPHEMERAL1, that controls petal senescence in Japanese morning glory. Plant J., 79, 1044–1051.

Shibuya, K., Shimizu, K., Yamada, T. and Ichimura, K. (2011) Expression of autophagy-associated *ATG8* genes during petal senescence in Japanese morning glory. J. Jap. Soc. Hort. Sci., 80, 89–95.

Shibuya, K., Watanabe, K. and Ono, M. (2018) CRISPR/Cas9-mediated mutagenesis of the *EPHEMERAL1* locus that regulates petal senescence in Japanese morning glory. Plant Physiol. Biochem., 131, 53–57.

Shibuya, K., Yamada, T. and Ichimura, K. (2009a) Autophagy regulates progression of programmed cell death during petal senescence in Japanese morning glory. Autophagy, 5, 546–547.

Shibuya, K., Yamada, T., Suzuki, T., Shimizu, K. and Ichimura, K. (2009b) InPSR26, a putative membrane protein, regulates programmed cell death during petal senescence in Japanese morning glory. Plant Physiol., 149, 816–824.

Shimoki, A., Tsugawa, S., Ohashi, K., Toda, M., Maeno, A., Sakamoto, T., Kimura, S., Nobusawa, T., Nagao, M., Nitasaka, E., Demura, T., Okada, K. and Takeda, S. (2021) Reduction in organ–organ friction is critical for corolla elongation in morning glory. *Commun*. Biol., 4, 285.

Shinozaki, Y., Tanabata, T., Ogiwara, I., Yamada, T. and Kanekatsu, M. (2011) Application of digital image analysis system for fine evaluation of varietal differences and the role of ethylene in visible petal senescence of morning glory. J. Plant Growth Regul., 30, 229–234.

Shinozaki, Y., Tanaka, R., Ono, H., Ogiwara, I., Kanekatsu, M., van Doorn, W.G. and Yamada, T. (2014) Length of the dark period affects flower opening and the expression of circadian-clock associated genes as well as xyloglucan endotransglucosylase/hydrolase genes in petals of morning glory (*Ipomoea nil*). Plant Cell Rep., 33, 1121–1131.

Shor, E., Skaliter, O., Sharon, E., Kitsberg, Y., Bednarczyk, D., Kerzner, S., Vainstein, D., Tabach, Y. and Vainstein, A. (2023) Developmental and temporal changes in petunia petal transcriptome reveal scent-repressing plant-specific RING-kinase-WD40 protein. Front. Plant Sci., 14, 1180899.

Stals, H. and Inzé, D. (2001) When plant cells decide to divide. Trends Plant Sci., 6, 359–364.

Stanke, M. and Waack, S. (2003) Gene prediction with a hidden Markov model and a new intron submodel. Bioinformatics, 19, Ii215–Ii225.

Strazzer, P., Verbree, B., Bliek, M., Koes, R. and Quattrocchio, F.M. (2023) The Amsterdam petunia germplasm collection: A tool in plant science. Front. Plant Sci., 14, 1129724.

Sun, L., Luo, H., Bu, D., Zhao, G., Yu, K., Zhang, C., Liu, Y., Chen, R. and Zhao, Y. (2013) Utilizing sequence intrinsic composition to classify protein-coding and long non-coding transcripts. Nucleic Acids Res., 41, e166.

Tanigaki, Y., Higashi, T., Takayama, K., Nagano, A.J., Honjo, M.N. and Fukuda, H. (2015) Transcriptome analysis of plant hormone-related tomato (*Solanum lycopersicum*) genes in a sunlight-type plant factory. PLoS One, 10.

Trunschke, J., Lunau, K., Pyke, G.H., Ren, Z.X. and Wang, H. (2021) Flower color evolution and the evidence of pollinator-mediated selection. Front. Plant Sci., 12, 617851.

van Doorn, W.G. and Kamdee, C. (2014) Flower opening and closure: an update. J. Exp. Bot., 65, 5749–5757.

van Doorn, W.G. and van Meeteren, U. (2003) Flower opening and closure: a review. J. Exp. Bot., 54, 1801–1812.

van Doorn, W.G. and Woltering, E.J. (2008) Physiology and molecular biology of petal senescence. J. Exp. Bot., 59, 453–480.

Verlinden, S. (2003) Changes in mineral nutrient concentrations in petunia corollas during development and senescence. HortSci., 38, 71–74.

Waki, T., Mameda, R., Nakano, T., Yamada, S., Terashita, M., Ito, K., Tenma, N., Li, Y., Fujino, N., Uno, K., Yamashita, S., Aoki, Y., Denessiouk, K., Kawai, Y., Sugawara, S., Saito, K., Yonekura-Sakakibara, K., Morita, Y., Hoshino, A., Takahashi, S. and Nakayama, T. (2020) A conserved strategy of chalcone isomerase-like protein to rectify promiscuous chalcone synthase specificity. Nat. Commun., 11, 870.

Wallace, H., Gel, Y. and Gastwirth, J. (2008) lawstat: An R Package for Law, Public Policy and Biostatistics. J. Stat. Softw., 28.

Wang, L., Li, Y., Jin, X., Liu, L., Dai, X., Liu, Y., Zhao, L., Zheng, P., Wang, X., Liu, Y., Lin, D. and Qin, Y. (2020) Floral transcriptomes reveal gene networks in pineapple floral growth and fruit development. Commun Biol, 3, 500.

Watanabe, K., Kobayashi, A., Endo, M., Sage-Ono, K., Toki, S. and Ono, M. (2017) CRISPR/Cas9-mediated mutagenesis of the *dihydroflavonol-4-reductase-B* (*DFR-B*) locus in the Japanese morning glory *Ipomoea (Pharbitis) nil*. Sci. Rep., 7, 10028.

Weller, J.L., Liew, L.C., Hecht, V.F.G., Rajandran, V., Laurie, R.E., Ridge, S., Wenden, B., Vander Schoor, J.K., Jaminon, O., Blassiau, C., Dalmais, M., Rameau, C., Bendahmane, A., Macknight, R.C. and Lejeune-Hénaut, I. (2012) A conserved molecular basis for photoperiod adaptation in two temperate legumes. Proc. Natl. Acad. Sci. USA, 109, 21158–21163.

Wozniak, N.J. and Sicard, A. (2018) Evolvability of flower geometry: Convergence in pollinator-driven morphological evolution of flowers. Semin. Cell Biol., 79, 3–15.

Wu, G., Anafi, R.C., Hughes, M.E., Kornacker, K. and Hogenesch, J.B. (2016) MetaCycle: an integrated R package to evaluate periodicity in large scale data. Bioinformatics, 32, 3351–3353.

Xu, W., Dubos, C. and Lepiniec, L. (2015) Transcriptional control of flavonoid biosynthesis by MYB–bHLH–WDR complexes. Trends Plant Sci., 20, 176–185.

Yamada, T., Ichimura, K., Kanekatsu, M. and van Doorn, W.G. (2009) Homologs of genes associated with programmed cell death in animal cells are differentially expressed during senescence of petals. Plant Cell Physiol., 50, 610–625.

Yamaguchi, T., Fukada-Tanaka, S., Inagaki, Y., Saito, N., Yonekura-Sakakibara, K., Tanaka, Y., Kusumi, T. and Iida, S. (2001) Genes encoding the vacuolar Na^+^/H^+^ exchanger and flower coloration. Plant Cell Physiol., 42, 451–461.

Yin, X., Lin, X., Liu, Y., Irfan, M., Chen, L. and Zhang, L. (2020) Integrated metabolic profiling and transcriptome analysis of pigment accumulation in diverse petal tissues in the lily cultivar ‘Vivian’. BMC Plant Biol, 20, 446.

Zagotta, M.T., Hicks, K.A., Jacobs, C.I., Young, J.C., Hangarter, R.P. and MeeksWagner, D.R. (1996) The Arabidopsis *ELF3* gene regulates vegetative photomorphogenesis and the photoperiodic induction of flowering. Plant J., 10, 691–702.

