## Supplementary Figures for "Transcriptomic dynamics of petal development in the one-day flower species, Japanese morning glory (*Ipomoea nil*)"

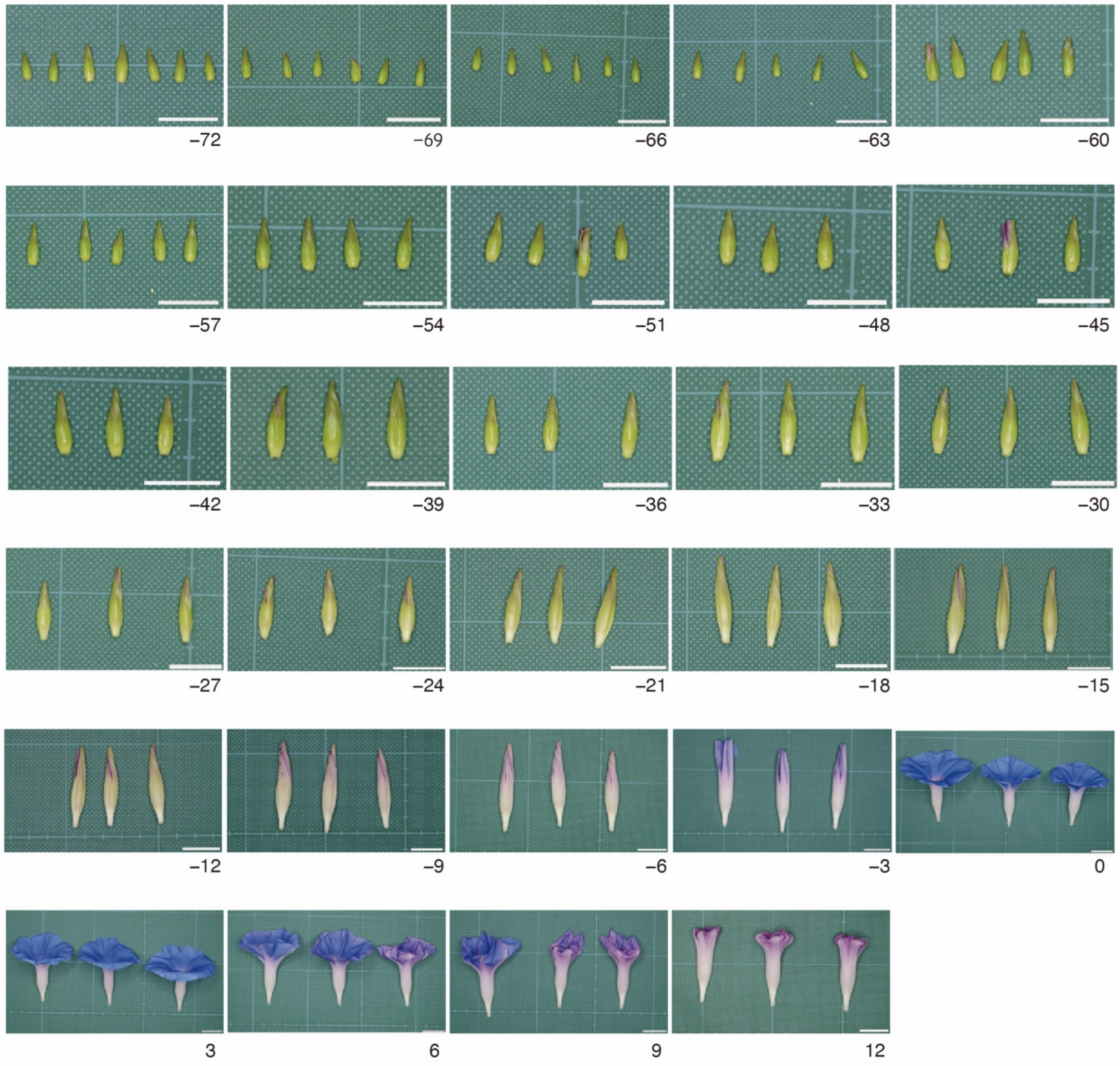


**Figure S1.** Petals used for RNA extraction.

Petals were sampled every 3 h, from 72 h before to 12 h after flower opening. The numbers below the images indicate the time relative to the flower opening. Scale bar, 10 mm.

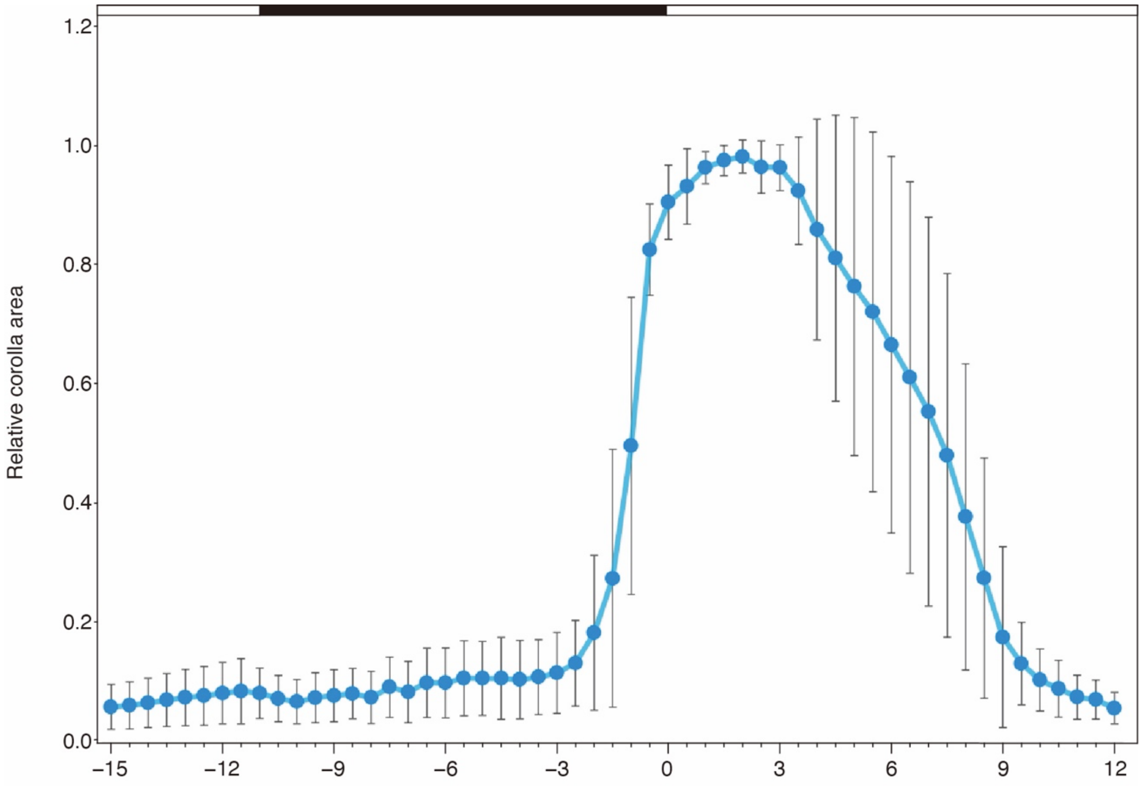


**Figure S2.** Measurement of relative corolla area.
The x-axis represents time to flower opening, while the y-axis shows the area ratio of the colored regions of the petals, expressed as the mean ± standard deviation (n = 10). The white and black bars at the top indicate light and dark conditions, respectively.

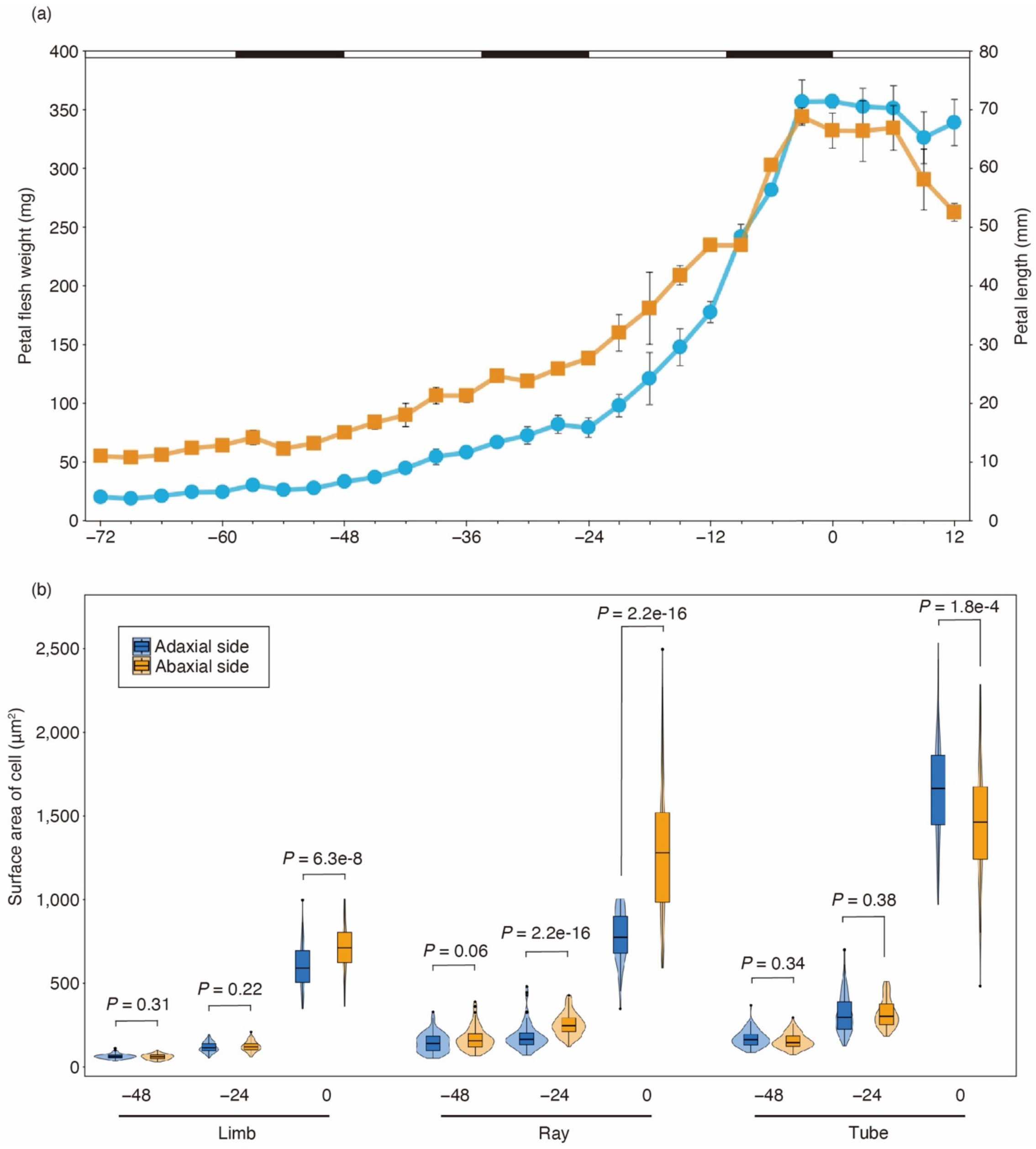


**Figure S3.** Temporal changes in the fresh weight and length of petals and measurement of the area of their epidermal cells.

(a) Fresh weight (orange) and length (blue) of petals from which RNA was extracted. The x-axis represents the time from flower opening. The white and black bars at the top indicate light and dark conditions, respectively. Values are expressed as the mean ± standard deviation, with 3–7 samples. The number of samples corresponds to the number of petals shown in Figure S1. (b) Area of epidermal cells. The x-axis represents the time from flower opening. Epidermal cells on the adaxial (blue) and abaxial (orange) sides of the limb, ray, and tube were observed using a scanning electron microscope, and their areas were calculated. The Brunner–Munzel test was used to compare the adaxial and abaxial sides ($n = 100$; $p < 0.05$).

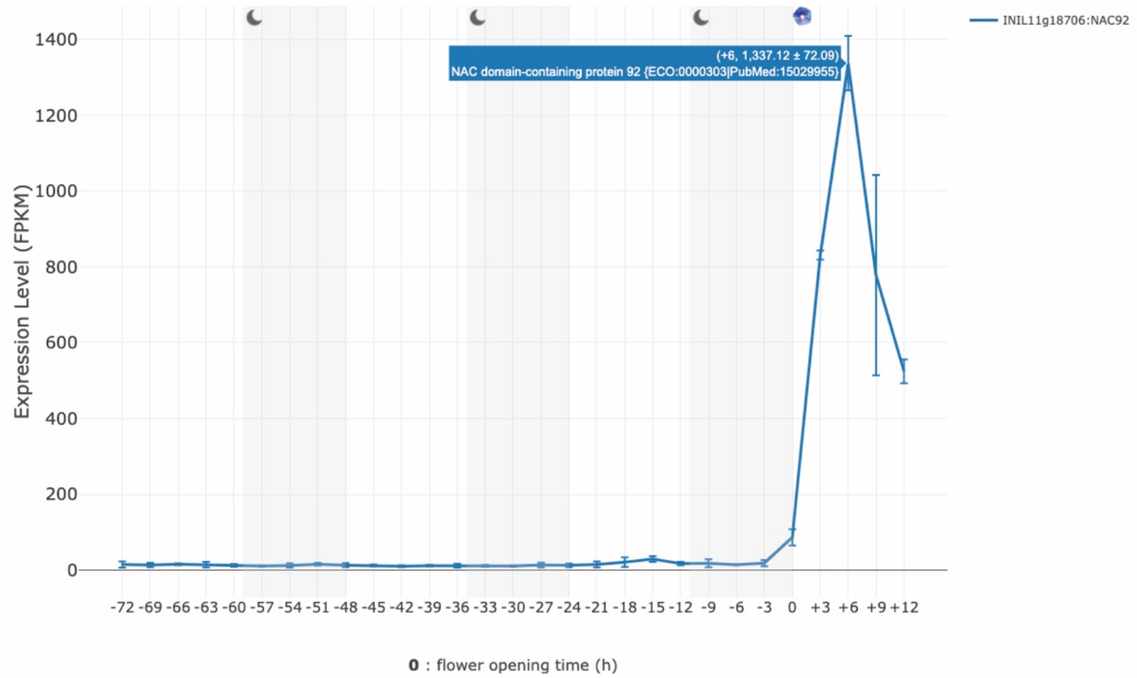


**Figure S4**. Expression pattern of the *EPH1* gene.
A screenshot from the database developed in this study showing the temporal gene expression levels in *Ipomoea nil* petals. The x-axis represents the time from flower opening (0), while the y-axis shows the mean ± standard deviation fragments per kilo base of transcript per million mapped fragments (FPKM) values for each time point (n = 3). The white and gray backgrounds indicate light and dark conditions, respectively.

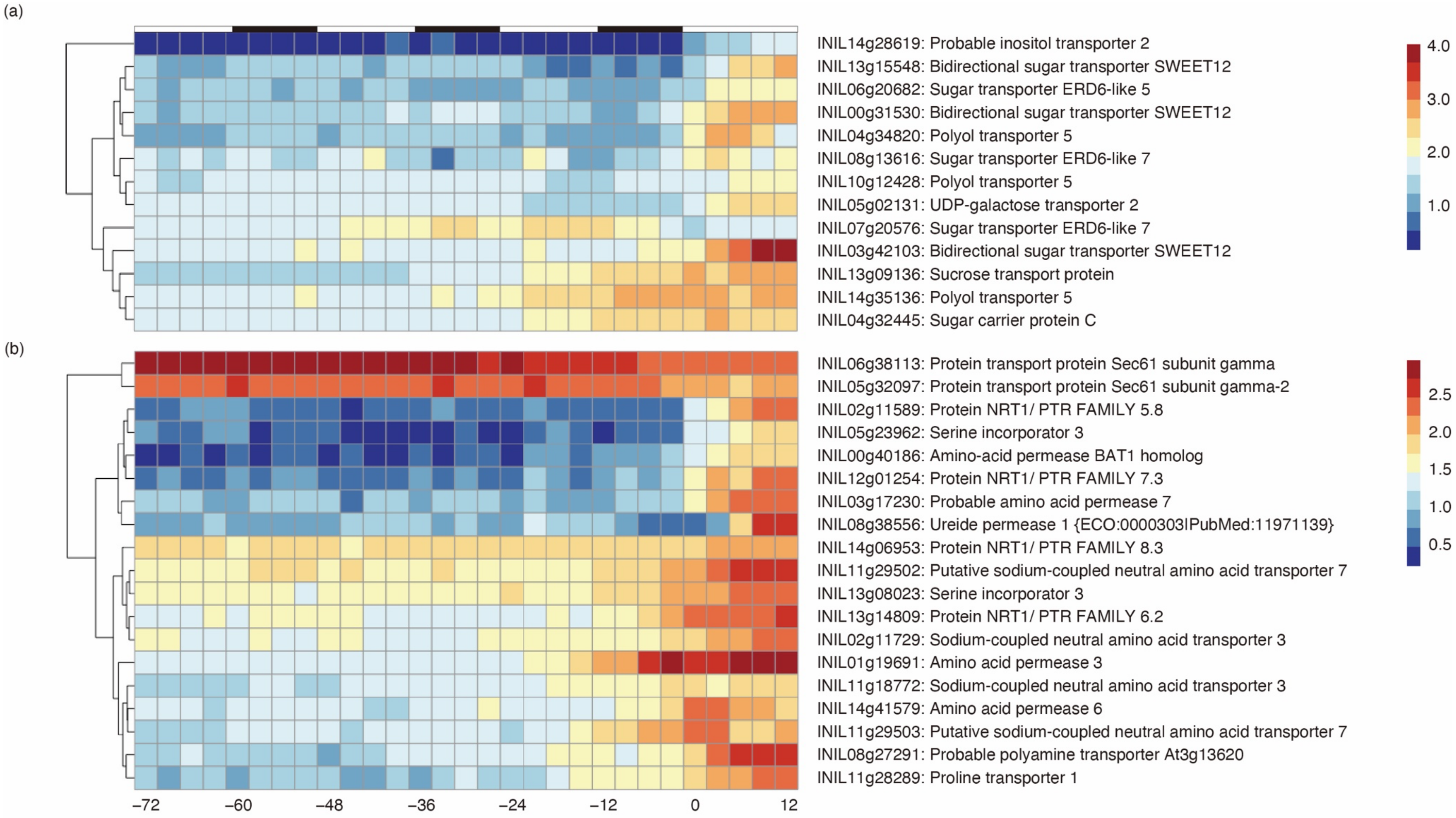
(a)
INIL14g28619: Probable inositol transporter 2
INIL13g15548: Bidirectional sugar transporter SWEET12
INIL06g20682: Sugar transporter ERD6-like 5
INIL00g31530: Bidirectional sugar transporter SWEET12
INIL04g34820: Polyol transporter 5
INIL08g13616: Sugar transporter ERD6-like 7
INIL10g12428: Polyol transporter 5
INIL05g02131: UDP-galactose transporter 2
INIL07g20576: Sugar transporter ERD6-like 7
INIL03g42103: Bidirectional sugar transporter SWEET12
INIL13g09136: Sucrose transport protein
INIL14g35136: Polyol transporter 5
INIL04g32445: Sugar carrier protein C
4.0
3.0
2.0
1.0
(b)
INIL06g38113: Protein transport protein Sec61 subunit gamma
INIL05g32097: Protein transport protein Sec61 subunit gamma-2
INIL02g11589: Protein NRT1/ PTR FAMILY 5.8
INIL05g23962: Serine incorporator 3
INIL00g40186: Amino-acid permease BAT1 homolog
INIL12g01254: Protein NRT1/ PTR FAMILY 7.3
INIL03g17230: Probable amino acid permease 7
INIL08g38556: Ureide permease 1 {ECO:0000303lPubMed:11971139}
INIL14g06953: Protein NRT1/ PTR FAMILY 8.3
INIL11g29502: Putative sodium-coupled neutral amino acid transporter 7
INIL13g08023: Serine incorporator 3
INIL13g14809: Protein NRT1/ PTR FAMILY 6.2
INIL02g11729: Sodium-coupled neutral amino acid transporter 3
INIL01g19691: Amino acid permease 3
INIL11g18772: Sodium-coupled neutral amino acid transporter 3
INIL14g41579: Amino acid permease 6
INIL11g29503: Putative sodium-coupled neutral amino acid transporter 7
INIL08g27291: Probable polyamine transporter At3g13620
INIL11g28289: Proline transporter 1
2.5
2.0
1.5
1.0
0.5
−72
−60
−48
−36
−24
−12
0
12

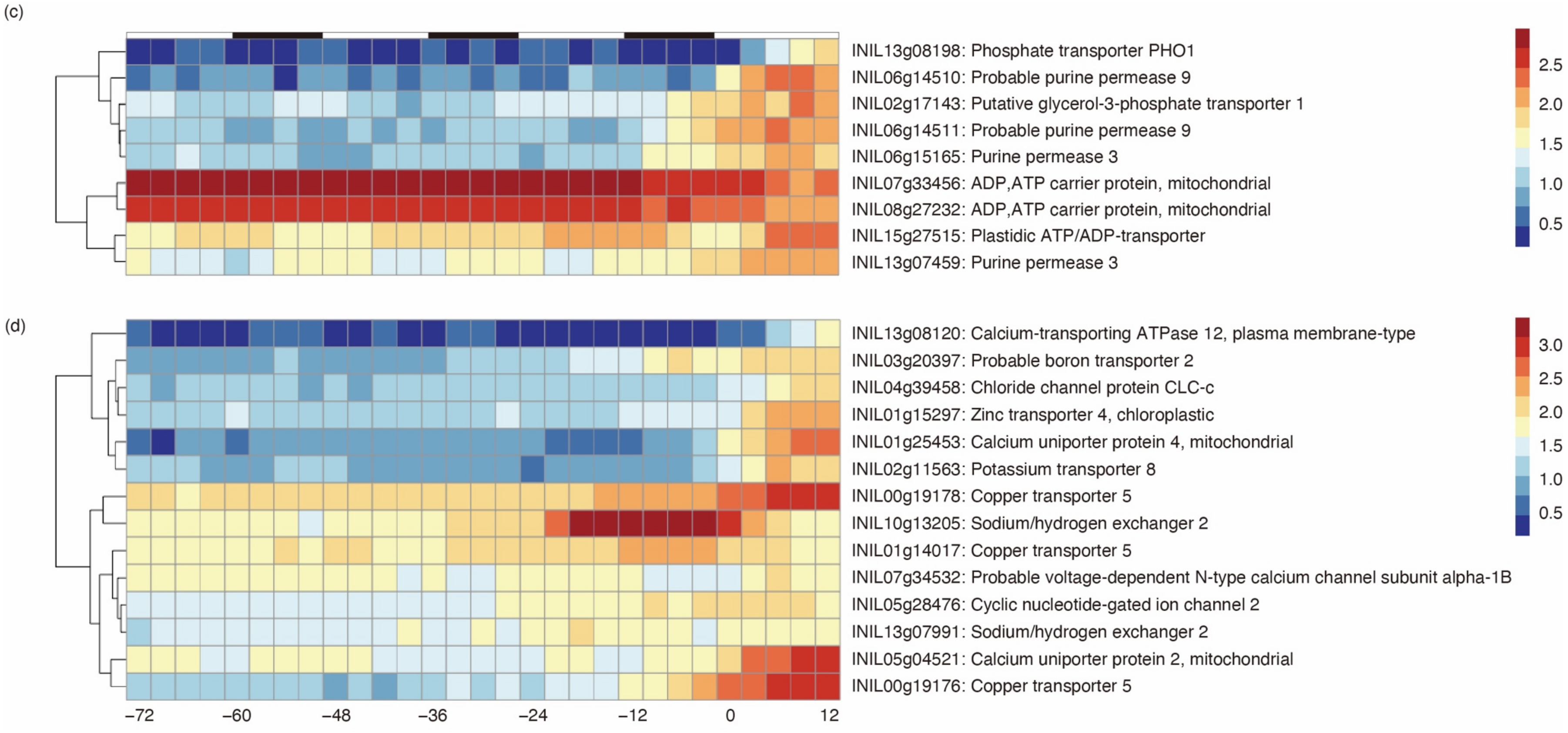


**Figure S5.** Heatmap analysis of transporter genes.
The temporal expression levels of genes associated with the seven transporter-related GO terms enriched at 12 h after flower opening (Figure 2 and Table S3) were visualized. Expression values are expressed as $\log_{10}$ (FPKM+1). The x-axis represents the time from flower opening (0). The white and black bars at the top represent light and dark conditions, respectively. Heatmaps are shown for transporter genes related to (a) sugars; (b) nitrogen (including amino acids); (c) phosphates (including nucleotides); and (d) inorganic ions.

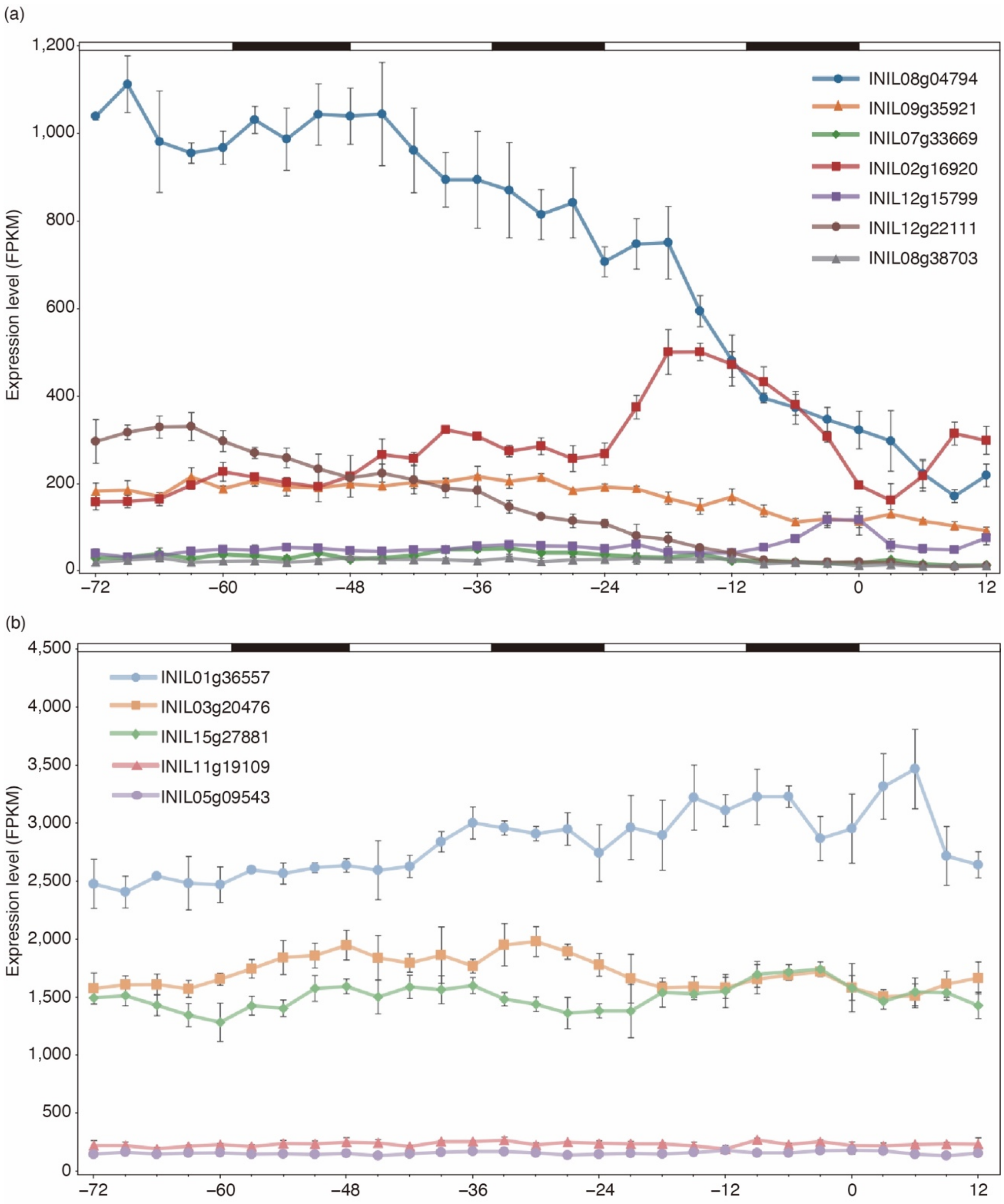


**Figure S6**. Expression patterns of internal control genes.
(a) Expression patterns of genes used as internal control genes in previous petal studies. (b) Expression patterns of the five genes with the lowest coefficient of variation among those with a mean fragments per kilo base of transcript per million mapped fragments (FPKM) value of 100 or higher. The x-axis represents the time from flower opening (0), while the y-axis shows the mean ± standard deviation FPKM values for each time point (n = 3). The white and black bars indicate light and dark conditions, respectively. Additional details, including references for each gene and their coefficients of variation, are provided in Table S4.

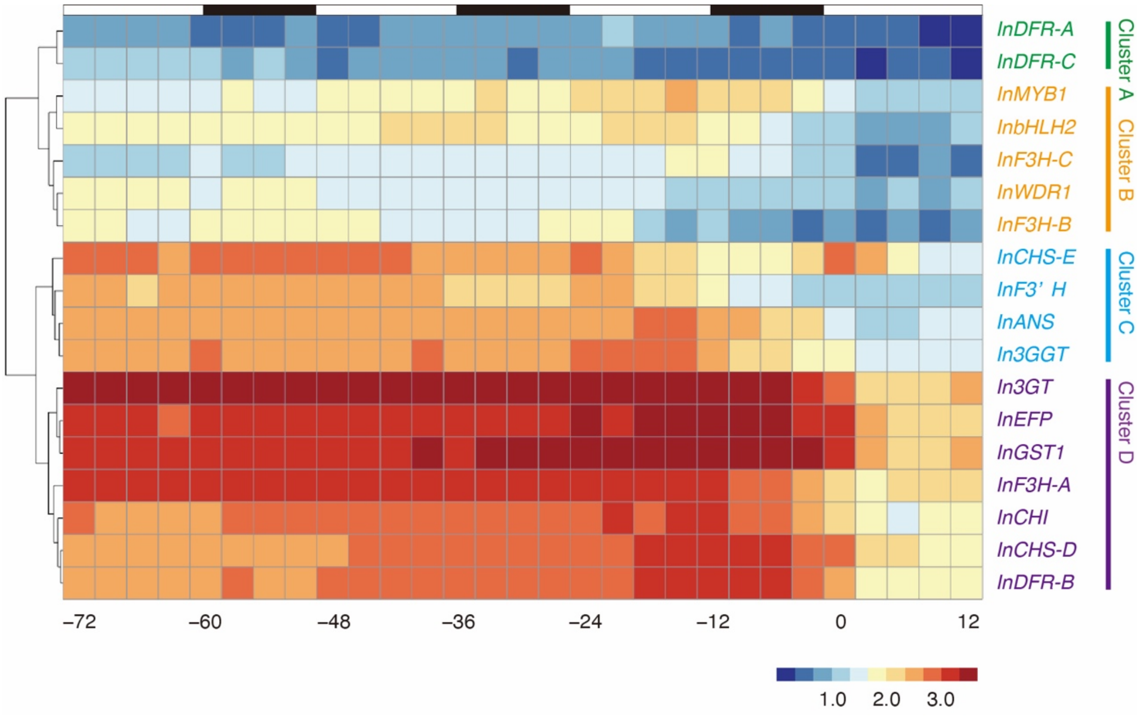


**Figure S7.** Heatmap analysis of the temporal expression levels of anthocyanin biosynthesis genes.

Table S15 lists the genes. Expression values are expressed as $\log_{10}$ (FPKM+1). The x-axis represents the time from flower opening (0), with the white and black bars at the top representing light and dark conditions, respectively. Gene names are color-coded by cluster, corresponding to the colors used for gene names in the heatmap in Figure 5a.

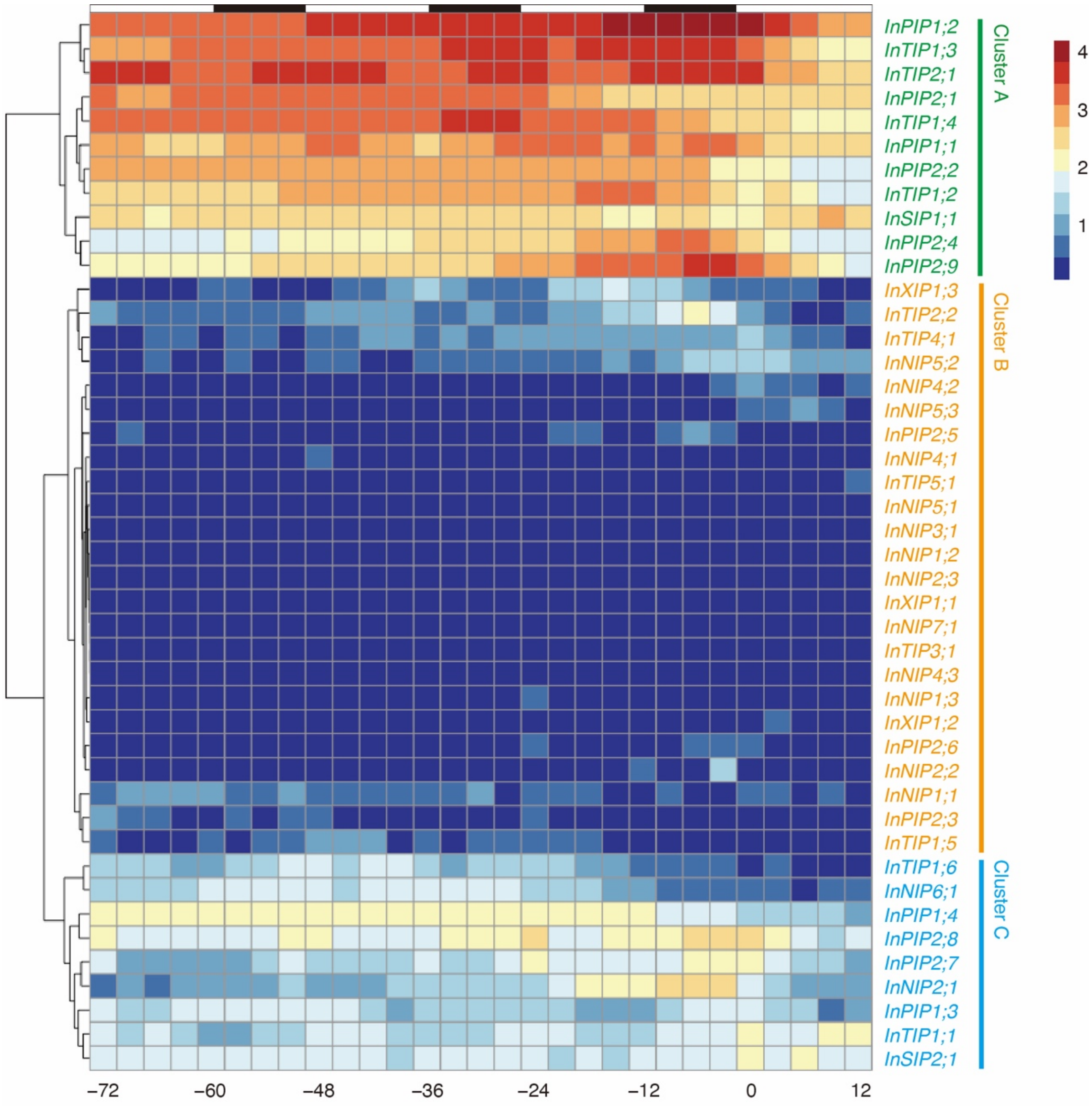


**Figure S8.** Heatmap analysis of the temporal expression levels of aquaporin genes. The list of the genes were obtained from Inden *et al*. (2023). Expression values are expressed as $\log_{10}$ (FPKM+1). The x-axis represents the time from the flower opening (0). The white and black bars at the top represent light and dark conditions, respectively. Gene names are color-coded by cluster, corresponding to the colors used for gene names in the heatmap in Figure 5b.

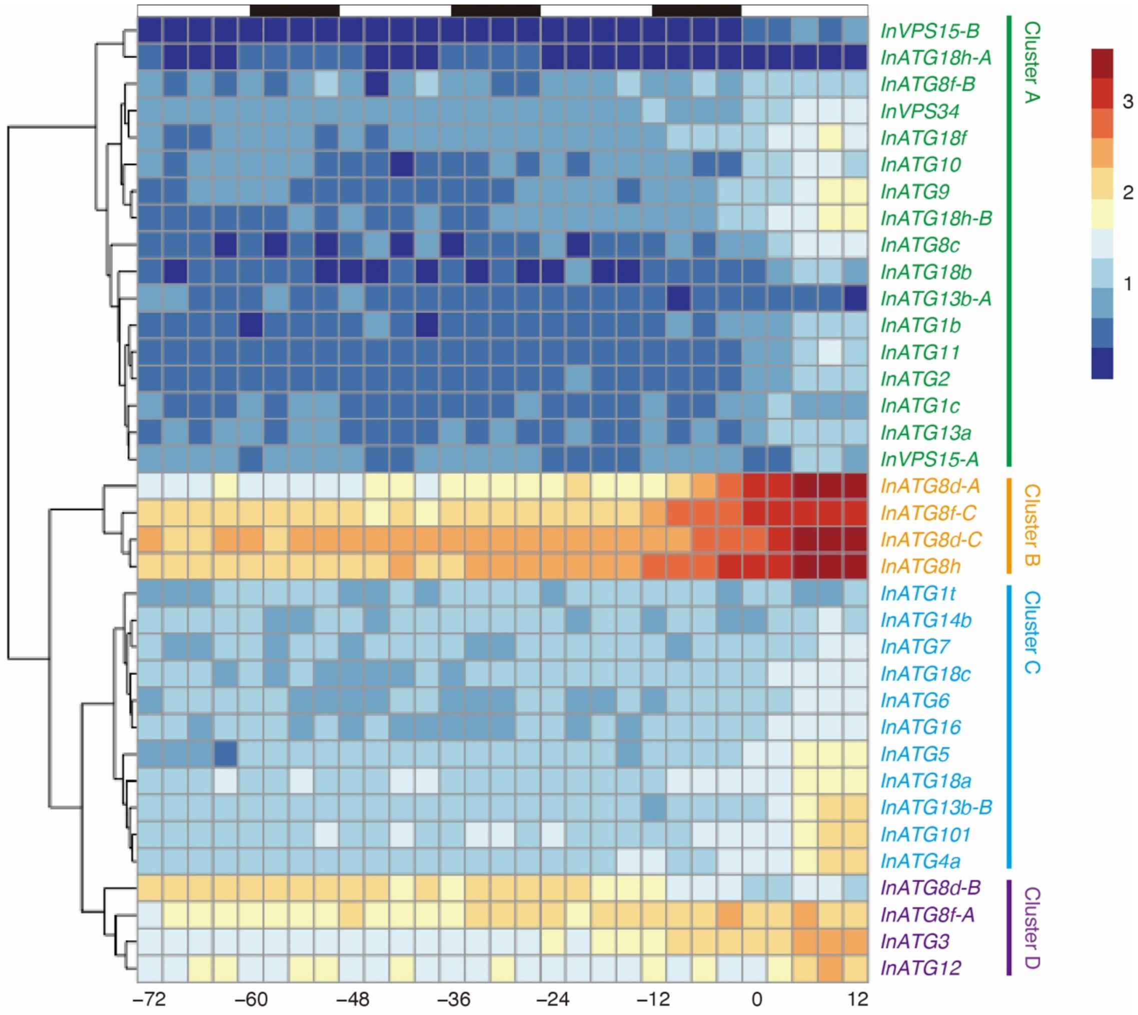


**Figure S9.** Heatmap analysis of the temporal expression levels of autophagy-related genes.

Table S18 provides the list of genes. Expression values are presented as $\log_{10}$ (FPKM+1). The x-axis represents the time from flower opening (0), with the white and black bars at the top representing light and dark conditions, respectively. Gene names are color-coded by cluster, corresponding to the colors used for gene names in the heatmap in Figure 5c.

### References

**Fukada-Tanaka, S., Hoshino, A., Hisatomi, Y., Habu, Y., Hasebe, M. and Iida, S.** (1997) Identification of new chalcone synthase genes for flower pigmentation in the Japanese and common morning glories. *Plant Cell Physiol.*, **38**, 754–758.

**Gutierrez, C.** (2009) The Arabidopsis cell division cycle. *Arabidopsis Book*, **7**, e0120.

**Hayama, R., Mizoguchi, T. and Coupland, G.** (2018) Differential effects of light-to-dark transitions on phase setting in circadian expression among clock-controlled genes in *Pharbitis nil*. *Plant Signal Behav.*, **13**, e1473686.

**Higuchi, Y., Sage-Ono, K., Sasaki, R., Ohtsuki, N., Hoshino, A., Iida, S., Kamada, H. and Ono, M.** (2011) Constitutive expression of the *GIGANTEA* ortholog affects circadian rhythms and suppresses one-shot induction of flowering in *Pharbitis nil*, a typical short-day plant. *Plant Cell Physiol.*, **52**, 638-650.

**Hoshino, A., Abe, Y., Saito, N., Inagaki, Y. and Iida, S.** (1997) The gene encoding flavanone 3-hydroxylase is expressed normally in the pale yellow flowers of the Japanese morning glory carrying the speckled mutation which produce neither flavonol nor anthocyanin but accumulate chalcone, aurone and flavanone. *Plant Cell Physiol.*, **38**, 970-974.

**Hoshino, A., Jayakumar, V., Nitasaka, E., Toyoda, A., Noguchi, H., Itoh, T., Shin-I, T., Minakuchi, Y., Koda, Y., Nagano, A.J., Yasugi, M., Honjo, M.N., Kudoh, H., Seki, M., Kamiya, A., Shiraki, T., Carninci, P., Asamizu, E., Nishide, H., Tanaka, S., Park, K.-I., Morita, Y., Yokoyama, K., Uchiyama, I., Tanaka, Y., Tabata, S., Shinozaki, K., Hayashizaki, Y., Kohara, Y., Suzuki, Y., Sugano, S., Fujiyama, A., Iida, S. and Sakakibara, Y.** (2016) Genome sequence and analysis of the Japanese morning glory *Ipomoea nil*. *Nat. Commun.*, **7**, 13295.

**Hoshino, A., Johzuka-Hisatomi, Y. and Iida, S.** (2001) Gene duplication and mobile genetic elements in the morning glories. *Gene*, **265**, 1–10.

**Hoshino, A., Morita, Y., Choi, J.D., Saito, N., Toki, K., Tanaka, Y. and Iida, S.** (2003) Spontaneous mutations of the flavonoid 3'-hydroxylase gene conferring reddish flowers in the three morning glory species. *Plant Cell Physiol.*, **44**, 990–1001.

**Hoshino, A., Park, K.I. and Iida, S.** (2009) Identification of *r* mutations conferring white flowers in the Japanese morning glory (*Ipomoea nil*). *J. Plant Res.*, **122**, 215–222.

**Inagaki, Y., Johzuka-Hisatomi, Y., Mori, T., Takahashi, S., Hayakawa, Y., Peyachoknagul, S., Ozeki, Y. and Iida, S.** (1999) Genomic organization of the genes encoding dihydroflavonol 4-reductase for flower pigmentation in the Japanese and common morning glories. *Gene*, **226**, 181-188.

**Inden, T., Hoshino, A., Otagaki, S., Matsumoto, S. and Shiratake, K.** (2023) Genome-wide analysis of aquaporins in Japanese morning glory (*Ipomoea nil*). *Plants*, **12**, 1511.

**Jaillon, O., Aury, J.-M., Noel, B., Policriti, A., Clepet, C., Casagrande, A., Choisne, N., Aubourg, S., Vitulo, N., Jubin, C., Vezzi, A., Legeai, F., Hugueney, P., Dasilva, C., Horner, D., Mica, E., Jublot, D., Poulain, J., Bruyère, C., Billault, A., Segurens, B., Gouyvenoux, M., Ugarte, E., Cattonaro, F., Anthouard, V., Vico, V., Del Fabbro, C., Alaux, M., Di Gaspero, G., Dumas, V., Felice, N., Paillard, S., Juman, I., Moroldo, M., Scalabrin, S., Canaguier, A., Le Clainche, I., Malacrida, G., Durand, E., Pesole, G., Laucou, V., Chatelet, P., Merdinoglu, D., Delledonne, M., Pezzotti, M., Lecharny, A., Scarpelli, C., Artiguenave, F., Pè, M.E., Valle, G., Morgante, M., Caboche, M., Adam-Blondon, A.-F., Weissenbach, J., Quétier, F., Wincker, P. and The French–Italian Public Consortium for Grapevine Genome, C.** (2007) The grapevine genome sequence suggests ancestral hexaploidization in major angiosperm phyla. *Nature*, **449**, 463-467.

**Kobayashi, K., Suzuki, T., Iwata, E., Magyar, Z., Bögre, L. and Ito, M.** (2015) MYB3Rs, plant homologs of Myb oncoproteins, control cell cycle-regulated transcription and form DREAM-like complexes. *Transcription*, **6**, 106-111.

**Lamesch, P., Berardini, T.Z., Li, D., Swarbreck, D., Wilks, C., Sasidharan, R., Muller, R., Dreher, K., Alexander, D.L., Garcia-Hernandez, M., Karthikeyan, A.S., Lee, C.H., Nelson, W.D., Ploetz, L., Singh, S., Wensel, A. and Huala, E.** (2012) The Arabidopsis Information Resource (TAIR): improved gene annotation and new tools. *Nucleic Acids Res*., **40**, D1202-D1210.

**Li, Y., Pi, M., Gao, Q., Liu, Z. and Kang, C.** (2019) Updated annotation of the wild strawberry *Fragaria vesca* V4 genome. *Hort. Res*., **6**, 61.

**McClung, C.R.** (2014) Wheels within wheels: new transcriptional feedback loops in the Arabidopsis circadian clock. *F1000Prime Rep*., **6**, 2.

**Morita, Y., Hoshino, A., Kikuchi, Y., Okuhara, H., Ono, E., Tanaka, Y., Fukui, Y., Saito, N., Nitasaka, E., Noguchi, H. and Iida, S.** (2005) Japanese morning glory *dusky* mutants displaying reddish-brown or purplish-gray flowers are deficient in a novel glycosylation enzyme for anthocyanin biosynthesis, UDP-glucose:anthocyanidin 3-*O*-glucoside-2″-*O*-glucosyltransferase, due to 4-bp insertions in the gene. *Plant J*., **42**, 353–363.

**Morita, Y., Ishiguro, K., Tanaka, Y., Iida, S. and Hoshino, A.** (2015) Spontaneous mutations of the UDP-glucose:flavonoid 3-*O*-glucosyltransferase gene confers pale- and dull-colored flowers in the Japanese and common morning glories. *Planta*, **242**, 575–587.

**Morita, Y., Saitoh, M., Hoshino, A., Nitasaka, E. and Iida, S.** (2006) Isolation of cDNAs for R2R3-MYB, bHLH and WDR transcriptional regulators and identification of *c* and *ca* mutations conferring white flowers in the Japanese morning glory. *Plant Cell Physiol*., **47**, 457–470.

**Morita, Y., Takagi, K., Fukuchi-Mizutani, M., Ishiguro, K., Tanaka, Y., Nitasaka, E., Nakayama, M., Saito, N., Kagami, T., Hoshino, A. and Iida, S.** (2014) A chalcone isomerase-like protein enhances flavonoid production and flower pigmentation. *Plant J*., **78**, 294–304.

**Nakamichi, N.** (2011) Molecular mechanisms underlying the *Arabidopsis* circadian clock. *Plant Cell Physiol*., **52**, 1709-1718.

**Norizuki, T., Kanazawa, T., Minamino, N., Tsukaya, H. and Ueda, T.** (2019) *Marchantia polymorpha*, a new model plant for autophagy studies. *Front Plant Sci*, **10**, 935.

**Ohnishi, M., Fukada-Tanaka, S., Hoshino, A., Takada, J., Inagaki, Y. and Iida, S.** (2005) Characterization of a novel $Na^+/H^+$ antiporter gene *InNHX2* and comparison of *InNHX2* with *InNHX1*, which is responsible for blue flower coloration by increasing the vacuolar pH in the Japanese morning glory. *Plant Cell Physiol*., **46**, 259–267.

**Ouyang, S., Zhu, W., Hamilton, J., Lin, H., Campbell, M., Childs, K., Thibaud-Nissen, F., Malek, R.L., Lee, Y., Zheng, L., Orvis, J., Haas, B., Wortman, J. and Buell, C.R.** (2007) The TIGR Rice Genome Annotation Resource: improvements and new features. *Nucleic Acids Res*., **35**, D883-D887.

**Park, K.I., Nitasaka, E. and Hoshino, A.** (2018) Anthocyanin mutants of Japanese and common morning glories exhibit normal proanthocyanidin accumulation in seed coats. *Plant Biotech*., **35**, 259–266.

**Sato, S., Nakamura, Y., Kaneko, T., Asamizu, E., Kato, T., Nakao, M., Sasamoto, S., Watanabe, A., Ono, A., Kawashima, K., Fujishiro, T., Katoh, M., Kohara, M., Kishida, Y., Minami, C., Nakayama, S., Nakazaki, N., Shimizu, Y., Shinpo, S., Takahashi, C., Wada, T., Yamada, M., Ohmido, N., Hayashi, M., Fukui, K., Baba, T., Nakamichi, T., Mori, H. and Tabata, S.** (2008) Genome structure of the Legume, *Lotus japonicus*. *DNA Res*, **15**, 227-239.

**Shemi, A., Ben-Dor, S. and Vardi, A.** (2015) Elucidating the composition and conservation of the autophagy pathway in photosynthetic eukaryotes. *Autophagy*, **11**, 701-715.

**Shibuya, K., Shimizu, K., Niki, T. and Ichimura, K.** (2014) Identification of a NAC transcription factor, EPHEMERAL1, that controls petal senescence in Japanese morning glory. *Plant J*., **79**, 1044-1051.

**Shibuya, K., Shimizu, K., Yamada, T. and Ichimura, K.** (2011) Expression of autophagy-associated *ATG8* genes during petal senescence in Japanese morning glory. *J. Jap. Soc. Hort. Sci.*, **80**, 89-95.

**Shibuya, K., Yamada, T., Suzuki, T., Shimizu, K. and Ichimura, K.** (2009) InPSR26, a putative membrane protein, regulates programmed cell death during petal senescence in Japanese morning glory. *Plant Physiol.*, **149**, 816-824.

**Shinozaki, Y., Tanaka, R., Ono, H., Ogiwara, I., Kanekatsu, M., van Doorn, W.G. and Yamada, T.** (2014a) Length of the dark period affects flower opening and the expression of circadian-clock associated genes as well as xyloglucan endotransglucosylase/hydrolase genes in petals of morning glory (*Ipomoea nil*). *Plant Cell Rep.*, **33**, 1121-1131.

**Shinozaki, Y., Tanaka, T., Ogiwara, I., Kanekatsu, M., van Doorn, W.G. and Yamada, T.** (2014b) Expression of an *AtNAP* gene homolog in senescing morning glory (*Ipomoea nil*) petals of two cultivars with a different flower life span. *J. Plant Physiol.*, **171**, 633-638.

**Watanabe, K., Oda-Yamamizo, C., Sage-Ono, K., Ohmiya, A. and Ono, M.** (2018) Alteration of flower colour in *Ipomoea nil t*hrough CRISPR/Cas9-mediated mutagenesis of *carotenoid cleavage dioxygenase 4*. *Transgenic Res.*, **27**, 25–38.

**Yamada, T., Ichimura, K., Kanekatsu, M. and van Doorn, W.G.** (2007) Gene expression in opening and senescing petals of morning glory (*Ipomoea nil*) flowers. *Plant Cell Rep.*, **26**, 823-835.

**Yamaguchi, T., Fukada-Tanaka, S., Inagaki, Y., Saito, N., Yonekura-Sakakibara, K., Tanaka, Y., Kusumi, T. and Iida, S.** (2001) Genes encoding the vacuolar $Na^+/H^+$ exchanger and flower coloration. *Plant Cell Physiol.*, **42**, 451–461.

**Zhou, Y., Zhang, Z., Bao, Z., Li, H., Lyu, Y., Zan, Y., Wu, Y., Cheng, L., Fang, Y., Wu, K., Zhang, J., Lyu, H., Lin, T., Gao, Q., Saha, S., Mueller, L., Fei, Z., Städler, T., Xu, S., Zhang, Z., Speed, D. and Huang, S.** (2022) Graph pangenome captures missing heritability and empowers tomato breeding. *Nature*, **606**, 527-534.
